# Two point mutations in protocadherin-1 disrupt Andes hantavirus recognition and afford protection against lethal infection

**DOI:** 10.1101/2022.07.19.500682

**Authors:** Megan M. Slough, Rong Li, Andrew S. Herbert, Gorka Lasso, Ana I. Kuehne, Russell R. Bakken, Stephanie R. Monticelli, Yanan Liu, Agnidipta Ghosh, Alicia M. Moreau, Xiankun Zeng, Steven C. Almo, John M. Dye, Rohit K. Jangra, Zhongde Wang, Kartik Chandran

**Author notes:** These authors contributed equally to this work (Megan M. Slough, Rong Li, Andrew S. Herbert). Senior author (Rohit K. Jangra, Zhongde Wang, Kartik Chandran).

## Abstract

Andes virus and Sin Nombre virus are the etiologic agents of severe hantavirus cardiopulmonary syndrome (HCPS) in the Americas for which no FDA-approved countermeasures are available. Protocadherin-1 (PCDH1), a cadherin-superfamily protein recently identified as a critical host factor for ANDV and SNV, represents a new antiviral target; however, its precise role remains to be elucidated. Here, we used computational and experimental approaches to delineate the binding surface of the ANDV glycoprotein complex on PCDH1’s first extracellular cadherin repeat domain. Strikingly, a single amino acid residue in this PCDH1 surface influenced the host species-specificity of SNV glycoprotein-PCDH1 interaction and cell entry. Mutation of this, and a neighboring residue, substantially protected Syrian hamsters from pulmonary disease and death caused by ANDV. We conclude that PCDH1 is a bona fide entry receptor for ANDV and SNV whose direct interaction with hantavirus glycoproteins could be targeted to develop new interventions against HCPS.

## Introduction

Rodent-borne orthohantaviruses (hereafter, hantaviruses) are segmented, negative-strand RNA viruses that have co-evolved with their rodent hosts over millions of years, by some estimates (1). Zoonotic transmission of some hantaviruses can cause two diseases in humans, hemorrhagic fever with renal syndrome (HFRS) and hantavirus cardiopulmonary syndrome (HCPS) (2). Both are associated with significant morbidity and mortality; case-fatality rates for HFRS and HCPS can approach 15% and 40%, respectively (3). No FDA-approved specific antivirals are available to treat HCPS or HFRS, and their development is challenged by the limited understanding of the hantavirus multiplication cycle.

Hantaviruses belonging to the ‘New World’ clade, Andes virus (ANDV) and Sin Nombre virus (SNV), are the major etiologic agents of HCPS in South and North America, respectively (2). ‘Old World’ hantaviruses found primarily in Asia and Europe, such as Hantaan virus (HTNV) and Seoul virus (SEOV), can cause HFRS in humans (2). We recently identified protocadherin-1 (PCDH1), a member of the non-clustered protocadherins in the cadherin superfamily (4,5), as a clade-specific entry host factor for New World hantaviruses, including ANDV and SNV, that directly engages the viral Gn/Gc glycoprotein complex (6,7), which mediates cell entry (8). PCDH1 is expressed in the airway epithelium, where it colocalizes with E-cadherin at apical cell-cell contact sites (9-11), and in vascular endothelial cells, the primary targets for hantavirus infection (12). *PCDH1* is a susceptibility gene for airway hyper-responsiveness and asthma (11,13,14), and its gene product appears to regulate airway epithelial barrier function through mechanisms that remain unclear (10).

PCDH1 is a Type I transmembrane protein comprising seven extracellular cadherin (EC) repeat domains, a ‘protocadherin domain’ that encompasses juxtamembrane and transmembrane sequences, and a long cytoplasmic tail (15,16). A soluble PCDH1 fragment, consisting of the first four EC repeat domains (EC1–4), crystallized as a head-to-tail homodimer making critical contacts between EC1 and EC4 (17). This suggests a mechanism by which PCDH1 molecules on neighboring cells form adhesive contacts (17). As reported previously, EC1 is necessary and sufficient for binding of PCDH1 by the New World hantavirus Gn/Gc, and PCDH1 is necessary for lethal ANDV infection in a Syrian hamster model of HCPS (6). However, the precise contacts at the PCDH1:Gn/Gc interface and the effects of PCDH1:Gn/Gc interaction on PCDH1 oligomerization (and vice versa) remain undefined.

Herein, we link the inter-species sequence variations in PCDH1 and mammalian host-specific receptor usage by hantaviruses to pinpoint sequences that could influence susceptibility to hantavirus infection. We then build on this information and combine structure-based prediction, comprehensive site-directed mutagenesis, and protein engineering to map Gn/Gc’s binding site in PCDH1 and shed light on the mode of hantavirus-PCDH1 recognition during viral entry.

## Results

### Murine lung microvascular endothelial cells are less susceptible to SNV Gn/Gc-dependent entry than their human counterparts

Natural infection by rodent-borne hantaviruses is proposed to be largely host-specific, suggesting the existence of molecular barriers to cross-species viral infection (3,18,19). Specifically, ANDV and SNV, whose reservoir hosts are the long-tailed pygmy rice rat (*Oryzomys longicaudatus*) and deer mouse (*Peromyscus maniculatus*), respectively, appear unable to infect the house mouse (*Mus musculus*) as no successful experimental infections of the latter have been reported. Further, mice engineered to lack a functional type I interferon response were reported to be highly susceptible to HTNV (20) and SEOV infection (21), but have not been shown to support ANDV or SNV infection. Thus, we postulated that there may be murine barriers to ANDV and SNV infection at the cellular level. To test this hypothesis, we infected primary human pulmonary microvascular endothelial cells (HPMECs) and primary murine lung microvascular endothelial cells (MLMECs) with replication-competent, recombinant vesicular stomatitis viruses (rVSVs) expressing ANDV (rVSV-ANDV-Gn/Gc) or SNV (rVSV-SNV-Gn/Gc) Gn/Gc. Mouse cells showed greatly (∼100-fold) reduced susceptibility to rVSV-SNV-Gn/Gc and slightly (∼5-fold) reduced susceptibility to rVSV-ANDV-Gn/Gc relative to the human cells (**Figure 1A**).

**Figure 1.**
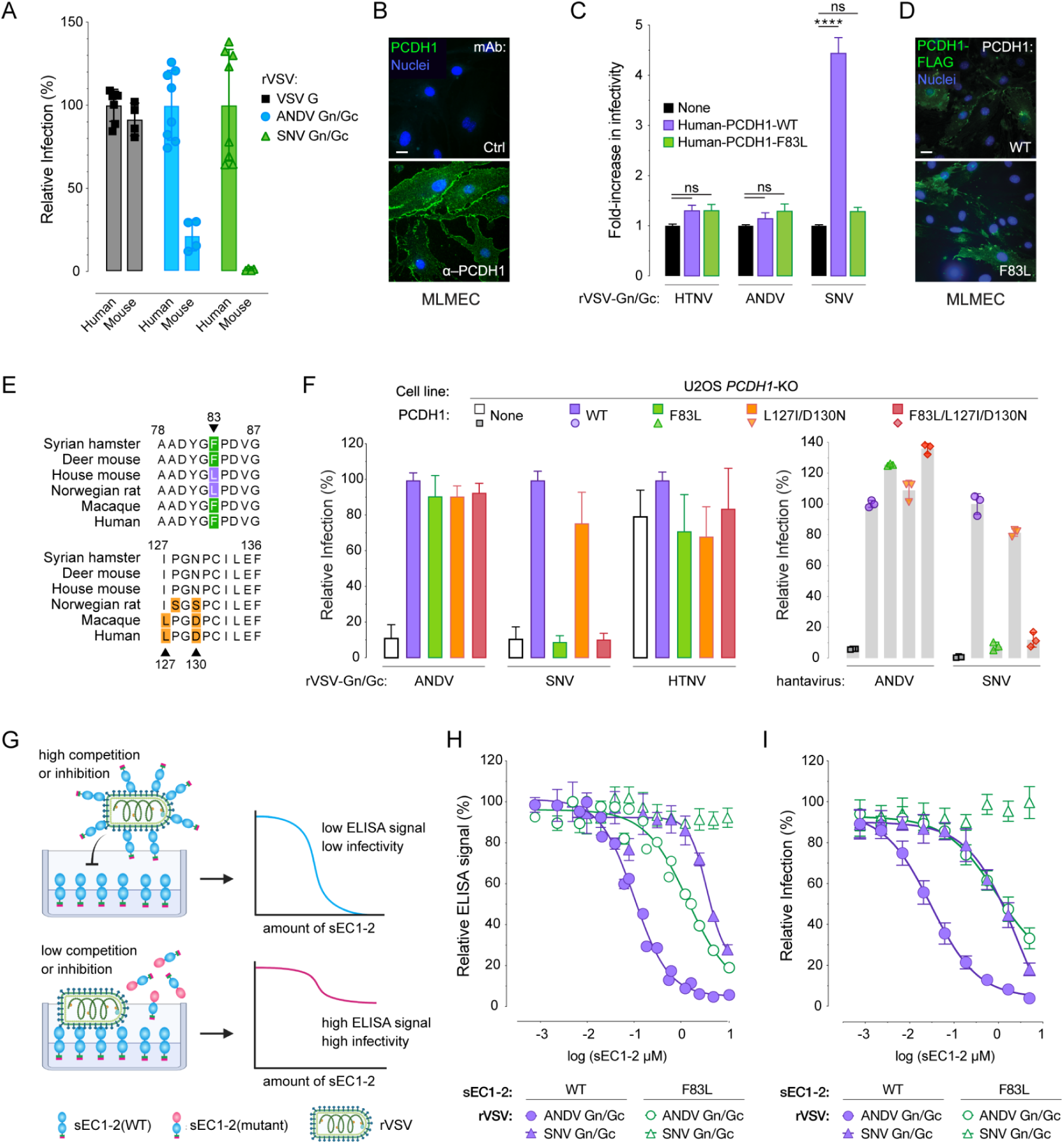
Residue F83 in PCDH1 is a key determinant of Sin Nombre virus infection. (A) Infectivity of rVSVs expressing G, ANDV Gn/Gc, or SNV Gn/Gc in human or mouse primary lung endothelial cells. Averages ± SD: two-four experiments, *n* = 4-8. (B) Expression of PCDH1 in primary mouse lung microvascular endothelial cells (MLMEC). Scale bar, 20 µm. Cells described in (B) were immunostained with PCDH1-specific monoclonal antibody (mAb)-3305 or a negative control mAb (Ctrl.) (C) Infectivity of rVSVs bearing HTNV-, ANDV-, or SNV-Gn/Gc in primary MLMECs expressing flag-tagged human WT or F83L-variant PCDH1. rVSV infectivities are expressed as fold change relative to that in non-complemented cells (set to one). Averages ± SD: three experiments, *n* = 35. Infectivities were compared by two-way ANOVA with Tukey’s correction for multiple comparisons; *ns* > 0.05, \*\*\*\**P* < 0.0001. (D) Cells described in (C) were immunostained with an anti-flag antibody. Scale bar, 20 µm. (E) Alignment of human PCDH1-EC1 amino acid sequences with those of a selection of rodent and primate species, highlighting residues that deviate from the consensus. Residues that specifically deviate from human EC1 are indicated [green, experimentally tested SNV-susceptible hosts; purple, experimentally unknown; orange, different residues in EC1 between human and other species with no known link to susceptibility]. Alignments generated by Clustal Omega. (F) The capacity of U2OS *PCDH1*-KO cells expressing the indicated PCDH1 variants to support hantavirus Gn/Gc-dependent entry. Cells were exposed to rVSVs bearing the indicated Gn/Gc proteins or authentic ANDV or SNV. “None” indicates no PCDH1 expression. Averages ± SD (rVSV): three experiments, *n* = 8. Averages ± SD (ANDV/SNV): one experiment, *n* = 2. (G) Diagram depicting competition ELISA and infection-inhibition assay comparing sEC1-2(WT) and sEC1-2(F83L) as competitive or inhibiting reagents. (H) Competition ELISA. rVSVs expressing ANDV or SNV Gn/Gc were pre-incubated with sEC1-2(WT) or sEC1-2(F83L) before added to sEC1-2(WT) coated ELISA plates. Averages ± SEM: two-three experiments, *n* = 4-7. (I) Infection-inhibition assay using sEC1-2(WT) and sEC1-2(F83L) to block infection of rVSVs bearing ANDV or SNV Gn/Gc on primary human endothelial cells (HUVECs). Averages ± SD: two experiments, *n* = 6. (sEC1-2, soluble extracellular cadherin domains 1 and 2).

### PCDH1 sequence variation at residue 83 influences endothelial cell susceptibility and SNV Gn/Gc:PCDH1 recognition

We postulated that differences in PCDH1 expression levels and/or sequence could account for the human-murine difference in viral entry into primary lung endothelial cells. However, immunostaining indicated that PCDH1 was abundantly expressed at the cell surface in mouse endothelial cells (**Figure 1B**). We found, instead, that ectopic expression of human PCDH1 in MLMEC enhanced infection by rVSV-SNV-Gn/Gc (**Figure 1C–D**), indicating that a molecular incompatibility between SNV Gn/Gc and murine PCDH1 is at least partially responsible for the entry block in murine endothelial cells.

We next examined the possibility that the murine ortholog of PCDH1 harbors amino acids that reduce its function as a hantavirus entry factor. Indeed, alignment of the EC1 domain sequences in PCDH1 from humans, non-human primates, and rodents revealed human-mouse sequence differences at three positions: F83L, L127I, and D130N (**Figure 1E**). To evaluate the effect of these EC1 residues on PCDH1’s receptor activity, we ectopically expressed EC1-‘murinized’ variants of human PCDH1 in *PCDH1-*knockout (KO) human osteosarcoma U2OS cells and tested their susceptibility to both, rVSVs bearing ANDV or SNV Gn/Gc and the authentic agents. Although all the PCDH1 variants expressed well and localized to the cell surface (**Figure S1A**), only the F83L substitution, alone or in combination with L127I and D130N, failed to restore cellular susceptibility to SNV entry and infection, suggesting F83L is determinative (**Figures 1F** **and S1B**). Moreover, over-expression of human PCDH1 bearing F83L failed to enhance rVSV-SNV-Gn/Gc infection in MLMECs (**Figures 1C–D**), supporting the conclusion that the human-murine sequence difference at PCDH1 position 83 renders murine endothelial cells less susceptible to SNV Gn/Gc-dependent entry.

Finally, we assessed the Gn/Gc binding activities of soluble, human PCDH1 comprising its first two EC domains (sEC1-2) (6) and bearing the human or murine residue at position 83 (wild-type “WT” or F83L, respectively). Pre-incubation of rVSV-ANDV-Gn/Gc with sEC1-2(WT), prevented the capture of rVSV particles by sEC1-2(WT) coated onto ELISA plates (**Figure 1H**) and inhibited cellular PCDH1-dependent viral entry (**Figure 1I**) in an sEC1-2 dose-dependent manner. As observed previously (6), SNV Gn/Gc was more resistant to sEC1-2 blockade than ANDV Gn/Gc in both of these assays, consistent with the lower affinity of the SNV Gn/Gc:PCDH1 interaction. By contrast, sEC1-2(F83L) displayed reduced activity against ANDV Gn/Gc and was devoid of activity against SNV Gn/Gc. Thus, F83L reduces PCDH1 binding to both ANDV and SNV Gn/Gc, but abolishes PCDH1 binding only to SNV Gn/Gc, concordant with its selective effect on entry and infection (**Figure 1F**). As shown previously, PCDH1 is dispensable for viral infection mediated by Old World hantavirus Gn/Gc (e.g., HTNV) (**Figure 1F**) (6,7). These findings, together with the conservation of F83 in PCDH1 orthologs from species known to be susceptible to New World hantaviruses (**Figure 1C**), provide evidence that Gn/Gc:PCDH1 recognition can influence the cellular host range of hantaviruses.

### Structure-based prediction of PCDH1:Gn/Gc interfacial residues

We postulated that F83 is a key contact residue within a larger Gn/Gc-binding surface in PCDH1. F83 is located in a disordered, membrane-distal loop in EC1 that is not resolved in the crystal structure of PCDH1 (encompassing EC1-4, PDB 6MGA) (17). We modeled multiple conformations of the missing loop, ranging from a more “closed” (buried) to a more “open” (solvent-accessible) conformation, which likely reflect the loop’s intrinsic flexibility (**Figure S2A**). In order to identify F83 neighboring residues that might also mediate Gn/Gc binding, we performed structure-based, interfacial prediction using the two most divergent modeled loop conformations, in which the F83 sidechain is either mostly buried or exposed to the solvent. We ran five complimentary algorithms to predict interfacial residues in the PCDH1 ectodomain and ranked predictions according to the number of supporting algorithms. The top 18 residues, predicted by at least four methods, were localized in the same face of the EC1 domain and included F83 (**Supplemental Item 1 and Figures S2B-C**). These residues were selected for further evaluation, and to capture additional, potential Gn/Gc-interacting residues that ranked lower in our analysis, we included 11 neighboring residues in our experimental studies of PCDH1:Gn/Gc binding (**Figures S2B-C**).

### Experimental mapping of the ANDV Gn/Gc:PCDH1 binding interface

We mutated the 29 selected EC1 residues to alanine (A) and/or residues with reversed charges or polarity (e.g., lysine (K) to aspartic acid (D) and alanine to serine (S); **Figure S2C**). sEC1-2 proteins bearing these point mutations were efficiently produced and largely exhibited similar electrophoretic mobilities (**Figure S3A**). Although three variants migrated anomalously during SDS-polyacrylamide gel electrophoresis (I140A, T142A, and R150A), PCDH1(I140A) eluted as a monodisperse peak with an apparent molecular weight resembling WT sEC1-2 (∼25-29K) in a size-exclusion column (6), indicating that it is largely monomeric in solution (**Figure S3B**).

The purified sEC1-2 mutants were screened for binding to ANDV Gn/Gc by competitive ELISA. Although some sEC1-2 mutations had no effect on PCDH1 binding (WT-like binders), others showed a minor or strong reduction in binding (intermediate and poor binders, respectively) (**Figure 2B**). Hierarchical clustering of the competition binding curves yielded two major clades separating the highly competitive mutants (where PCDH1 binding remains unaffected) from the poorly competitive ones (mutations that impair PCDH1 binding) (**Figure 2C**). Within each clade, subgroups that displayed intermediate activity or had mild reduction in binding could also be defined. Similar results were obtained in the infection-inhibition assay (**Figures 3A–B**), and a direct comparison of mutation effects in each assay showed that they were highly concordant (**Figure 3C**). Altogether, we found 11 EC1 residues whose mutation strongly impairs PCDH1 binding to ANDV Gn/Gc, 10 of which were predicted as interfacial by at least 4/5 algorithms and one (L152) was predicted with lower consent (3/5 algorithms) (**Figures S2B-C and 3C**). Interestingly, and in accordance with our mapping results, mutation of three residues identified as poor binders in this analysis, reduced recognition by a PCDH1-specific, monoclonal antibody previously shown to block PCDH1-Gn/Gc binding (6), with the strongest effect being observed for D85 (**Figure S4**). We have thus employed a combination of computational and experimental studies to uncover the ANDV Gn/Gc-binding surface in PCDH1 EC1. This surface comprises at least 11 residues and is centered around a flexible loop containing a residue (F83) that influences cellular susceptibility to viral entry in a host species-dependent manner.

**Figure 2.**
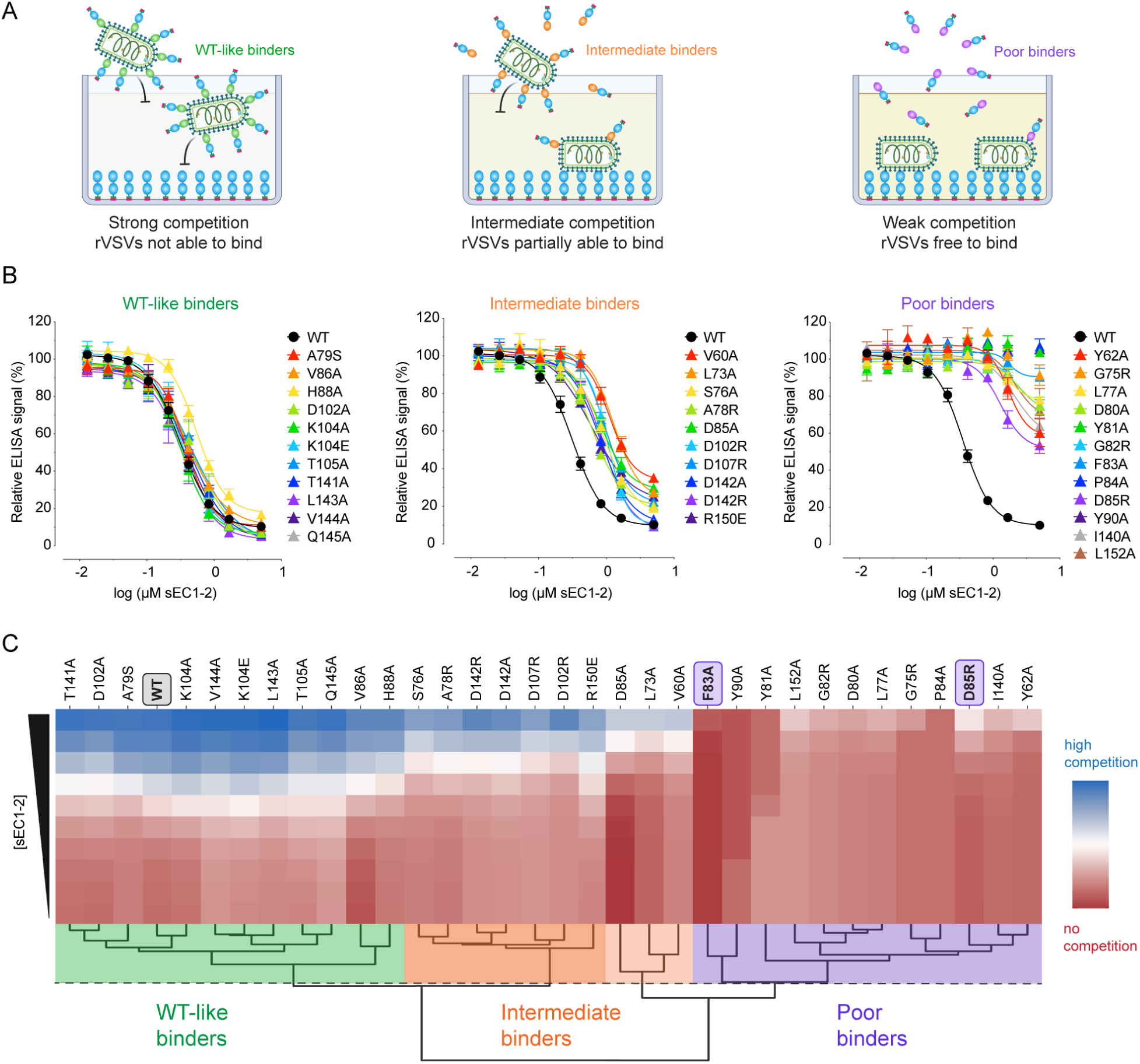
Binding capacity of mutant sEC1-2 to ANDV Gn/Gc. (A) Diagram of competition ELISA depicting three different competition outcomes of mutant sEC1-2 proteins’ capacity to block rVSV-ANDV-Gn/Gc binding to sEC1-2(WT) coated wells. (B) Competition ELISA using WT and mutant sEC1-2 as competitive reagents to the binding of rVSV-ANDV-Gn/Gc to sEC1-2(WT) coated wells. Averages ± SEM: three to four experiments, *n* = 6-8. (C) Hierarchical clustering of WT and mutant sEC1-2 generated from sigmoidal curves of the competition ELISA data in (A). The dendrogram shows four clusters, separating the poor binders from the WT-like binders. The dotted line denotes the height at which the dendrogram is cut representing varying degrees of binding strength; WT-like binders (green), intermediate binders (WT-like-intermediate binders in dark orange, intermediate-poor binders in light orange), and poor binders (purple).

**Figure 3.**
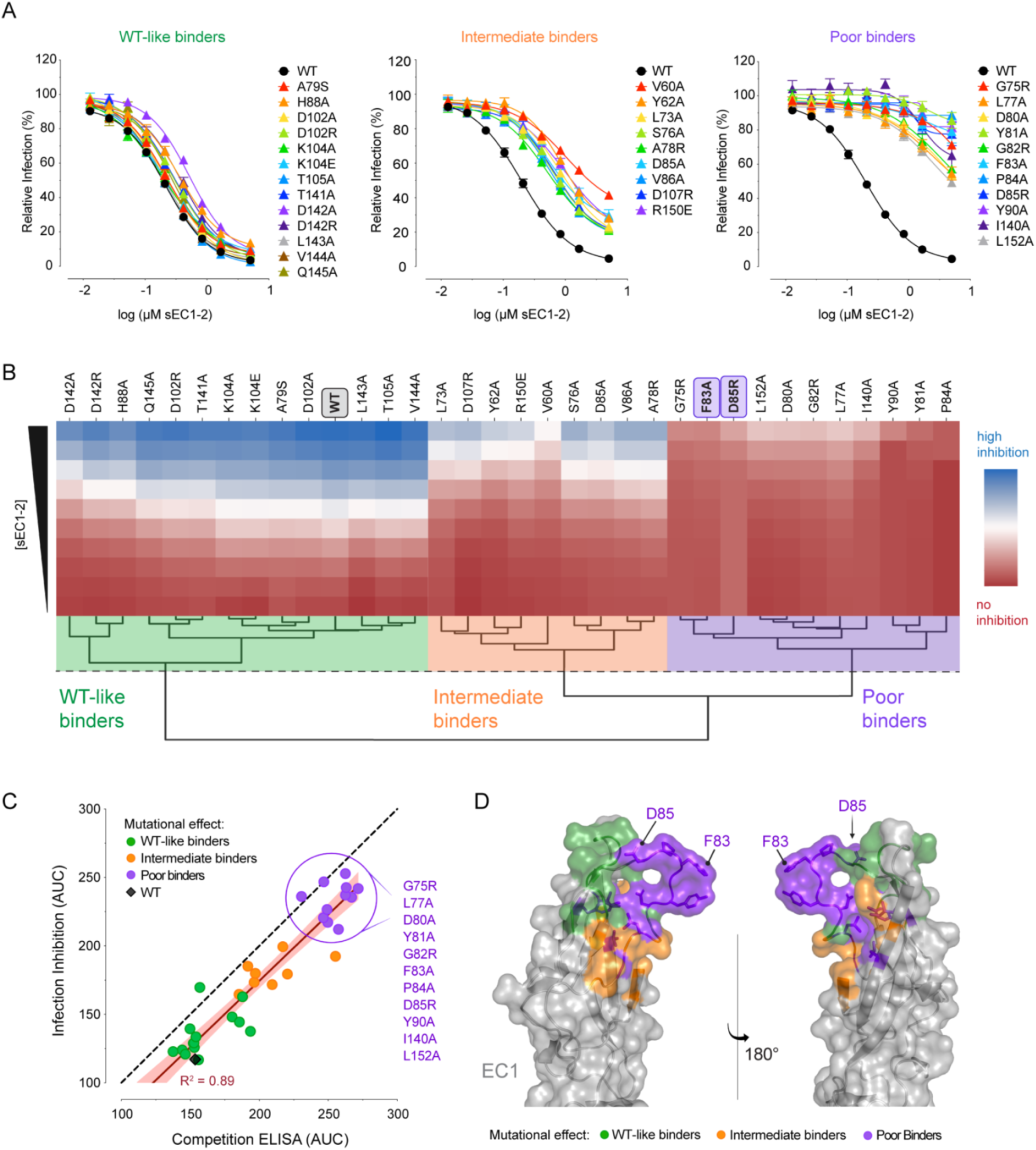
Inhibition of ANDV Gn/Gc-mediated infection by mutant sEC1-2. (A) WT and mutant sEC1-2 were tested on their ability to block rVSV-ANDV-Gn/Gc entry in primary human endothelial cells (HUVECs). Averages ± SEM: three experiments, *n* = 5-7. (B) Hierarchical clustering of WT and mutant sEC1-2 generated from sigmoidal curves of the infection-inhibition assay in (A). The dotted line denotes the height at which the dendrogram is cut to obtain three clusters representing varying degrees of inhibition of ANDV Gn/Gc-initiated infection; WT-like inhibition (green), intermediate inhibition (orange), and poor inhibition (purple). (C) Area under the curve (AUC) for the binding activity of WT and mutant sEC1-2 to rVSV-ANDV-Gn/Gc, as determined by competition ELISA (see Figure 2), plotted against AUC values as determined by rVSV-ANDV-Gn/Gc infection-inhibition assay (A). The red line denotes the R squared value. A list of the sEC1-2 mutants that are classified as poor binders are listed to the right. (D) EC1 crystal structure in the “open conformation” displaying mutated residues representing three degrees of binding strength to ANDV Gn/Gc and inhibition of rVSV-ANDV-Gn/Gc infection. sEC1-2 mutants that bind and inhibit similarly to WT (WT-like binders), green; sEC1-2 mutants that display a mild reduction in binding and inhibition (intermediate binders), orange, and sEC1-2 mutants that display a strong reduction in binding and inhibition (poor binders), purple. Structure adapted from PDB 6MGA.

### Key Gn/Gc-binding residues in PCDH1 are required for hantavirus entry and infection

We identified a PCDH1 EC1 surface patch comprising 11 residues that impair ANDV Gn/Gc binding. Next, we evaluated the roles of two individual residues—F83, which influences cellular susceptibility to viral entry (**Figure 1**), and D85, which is a key residue in the epitope of a mAb that blocks PCDH1 binding (**Figure S4**). We first generated *PCDH1-*knockout (KO) U2OS cell lines stably expressing full-length PCDH1 clones bearing mutations in F83 and D85, or at an adjacent residue not implicated in ANDV Gn/Gc binding, V86. All of the PCDH1 variants resembled WT in expression level and localization at the cell surface (**Figure S5**). We evaluated the cells’ susceptibility to hantavirus Gn/Gc-dependent infection and found that mutating either F83 or D85 was sufficient to block infection by both rVSV-SNV-Gn/Gc (**Figure 4A**) and authentic SNV (**Figure 4B**). By contrast, and consistent with the higher affinity of the ANDV Gn/Gc:PCDH1 interaction, the combination of both mutations was necessary to reduce ANDV Gn/Gc-mediated infection (**Figure 4A**). The V86A PCDH1 variant had no effect on entry by either viral glycoprotein, consistent with its dispensability for PCDH1 recognition by Gn/Gc (**Figure 2B**). These findings support that the identified Gn/Gc-binding surface in PCDH1 is required for cell entry by two virulent New World hantaviruses.

**Figure 4.**
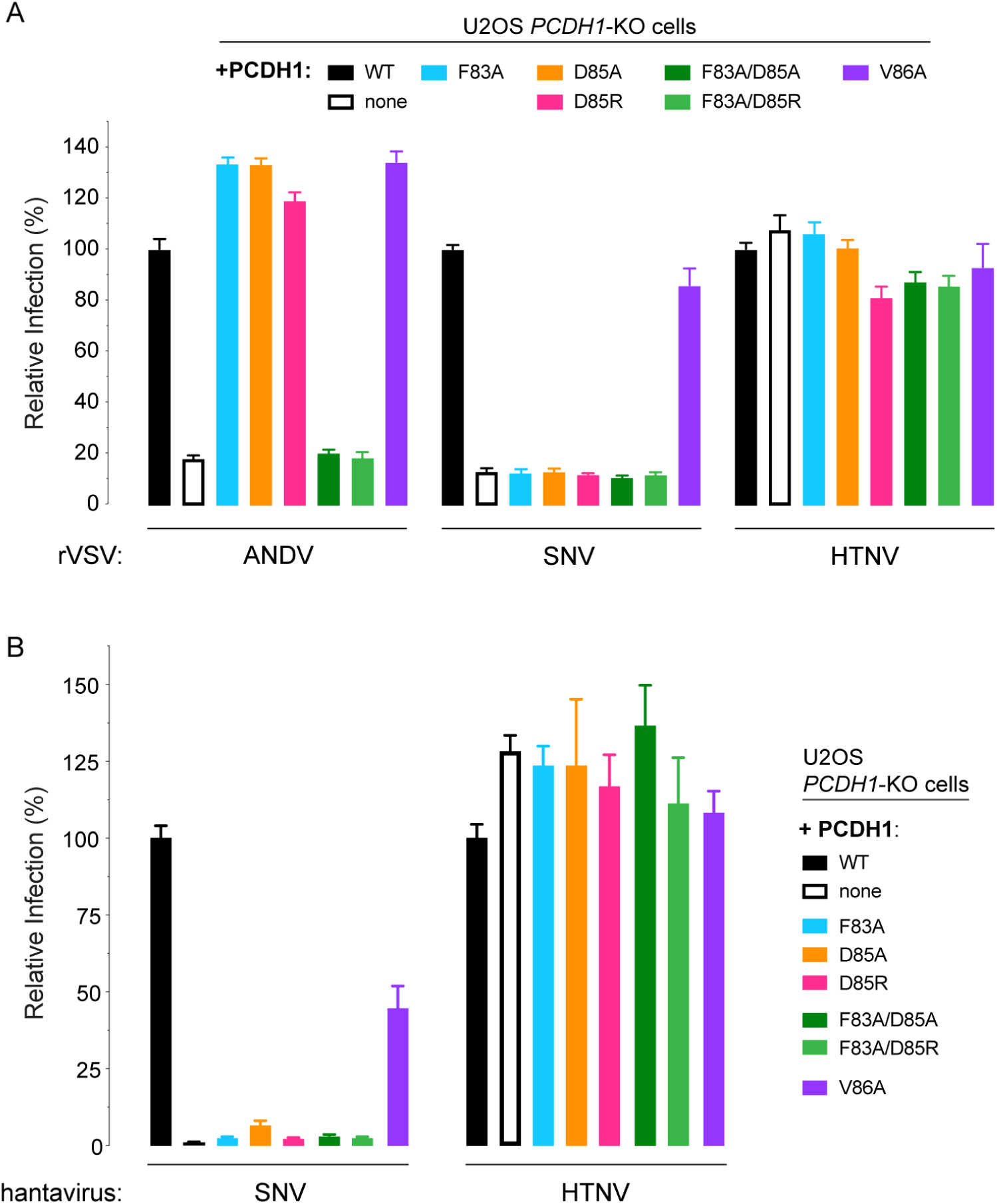
Two key amino acids in PCDH1 mediate entry for SNV and ANDV. (A) Relative infectivity of rVSVs bearing ANDV-, SNV-, or HTNV-Gn/Gc on U2OS *PCDH1-*KO cells complemented with WT or mutant PCDH1. Averages ± SEM: three to four experiments, *n* = 8-12. (B) Relative infectivity of authentic SNV or HTNV on the cell lines described in (A). Averages ± SEM: two to three experiments, *n* = 6-9.

### PCDH1 homodimerization is dispensable for Gn/Gc recognition

Previous work has shown that the PCDH1 ectodomain can form head-to-tail homodimers, driven largely by intermolecular interactions between the EC1 and EC4 domains (17) (**Figure 5A**). Although the EC4 and Gn/Gc contact sites in EC1 do not appear to overlap, we considered the possibility that EC1 dimerization may nevertheless impact Gn/Gc recognition. Accordingly, we tested the effects of mutations at EC1 position E137 (**Figure S6A**), responsible for driving the EC1:EC4 interaction (17), on ANDV Gn/Gc binding. These mutants resembled WT in their capacity to compete for ANDV Gn/Gc binding (**Figure 5B**). To further examine the effect of PCDH1 dimerization on its hantavirus receptor activity, we stably expressed PCDH1 lacking the EC4 domain in *PCDH1*-KO U2OS cells (**Figure S6B**) and tested their susceptibility to Gn/Gc-mediated infection (**Figure 5C**). This “monomer-only” PCDH1(ΔEC4) supported ANDV and SNV Gn/Gc-dependent entry similar to WT PCDH1.

**Figure 5.**
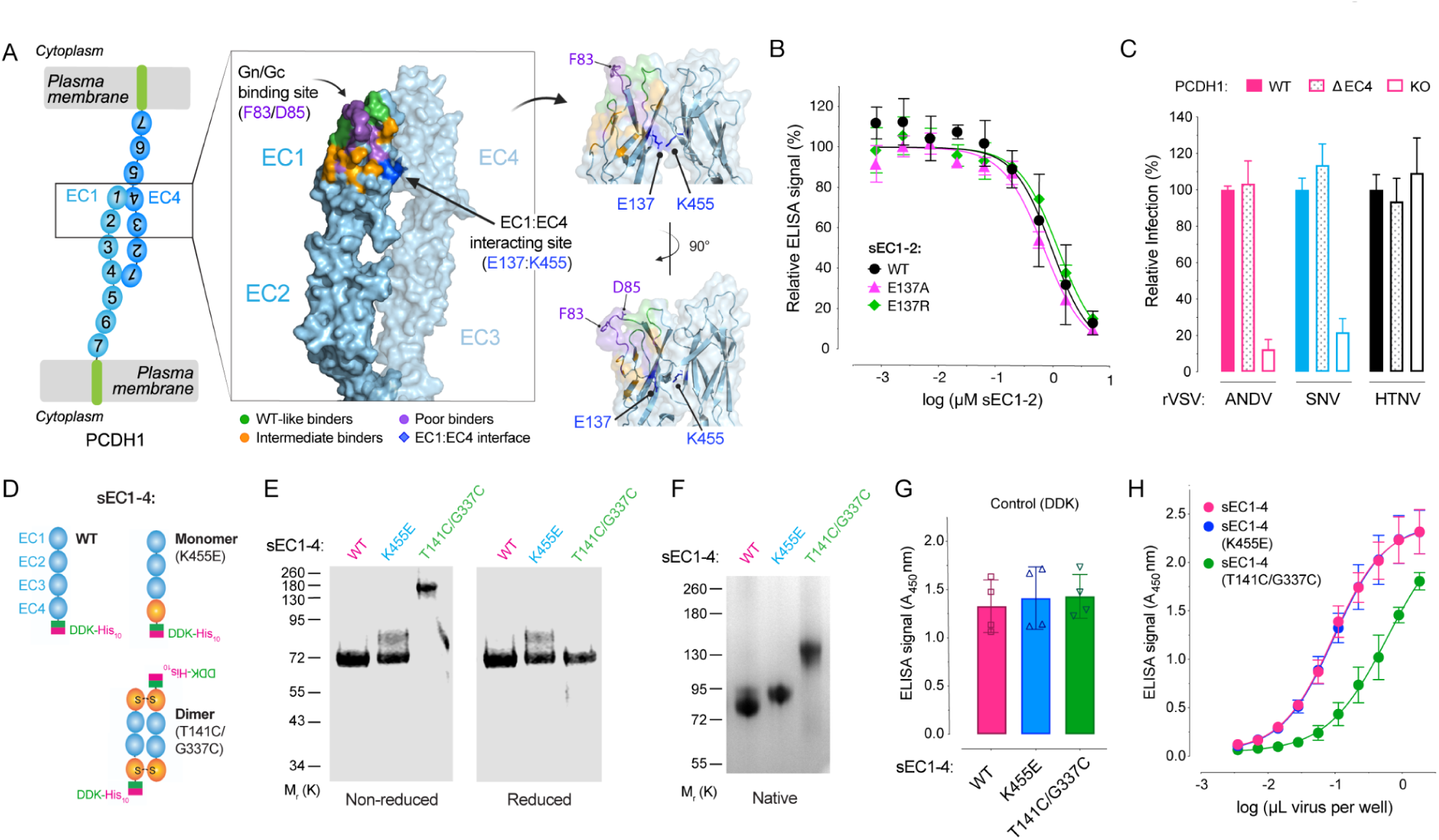
Monomeric and dimeric PCDH1 provide suitable entry receptors for ANDV. (A) Crystal structure of the proposed anti-parallel EC1-4 trans-dimer. Structure is in the “open conformation” displaying mutated residues representing three degrees of binding strength to ANDV Gn/Gc. sEC1-2 mutants that bind similarly to WT (WT-like binders), green; sEC1-2 mutants that display a mild reduction in binding (intermediate binders), orange, and sEC1-2 mutants that display a strong reduction in binding (poor binders), purple. The binding site including residues F83 and D85, and the EC1:EC4 adhesive interface are indicated. A closer view of the EC1:EC4 binding interface is shown to the right. Structure adapted from PDB 6MGA. (B) Competition ELISA using WT and mutant sEC1-2 as competitive reagents to the binding of rVSV-ANDV-Gn/Gc to WT sEC1-2 coated wells. Averages ± SD: two experiments, *n* = 3-4. (C) Relative infectivity of rVSVs bearing ANDV-, SNV-, or HTNV-Gn/Gc on U2OS *PCDH1-*KO cells complemented with WT or ΔEC4 PCDH1. (Averages ± SD: three experiments, *n* = 9) (D) Schematic representation of WT or mutant sEC1-4 proteins forming monomers or dimers. (E) Non-reduced and reduced purified WT and mutant sEC1-4 were separated on an SDS-polyacrylamide gel and visualized by Coomassie Brilliant Blue staining. M_r_, relative molecular weight (K denotes x 1,000). (F) Non-reduced samples in (E) were run on a native-polyacrylamide gel electrophoresis and visualized as in (E). (G) ELISA detecting Flag-tagged WT and mutant sEC1-4 coated plates, using an anti-Flag-HRP antibody. Done in parallel with (H). Averages ± SD: two experiments, *n* = 4. (H) Capacity of rVSVs bearing ANDV Gn/Gc to bind to WT or mutant sEC1-4 coated plates. Averages ± SD: two experiments, *n* = 4. (sEC1-4, soluble extracellular cadherin domains 1-4)

PCDH1 interaction and homodimerization *in trans* is proposed to regulate cell adhesion between neighboring cells, suggesting that both monomers and dimers co-exist at the cell surface (17). Although our preceding results indicated that hantaviruses could efficiently recognize and use PCDH1 monomers, they did not exclude the possibility that PCDH1 dimers may also provide suitable entry receptors. To test this, we engineered sEC1-4 variants that are intrinsically either monomers or dimers (**Figure 5D**). Specifically, we generated obligate monomers by introducing the K455E mutation in EC4 to abrogate its salt bridge with E137 in EC1, disrupting the EC1:EC4 interaction (17). To generate obligate dimers, we engineered a sEC1-4 variant in which structurally apposed residues, T141 and G337 in EC1 and EC4, respectively, were mutated to C to afford intersubunit disulfide formation. sEC1-4(T141C/G337C) displayed shifts in electrophoretic mobility relative to sEC1-4 (WT) and sEC1-4(K455E) monomer at nonreducing conditions in denaturing (**Figure 5E**) and native polyacrylamide gels (**Figure 5F**), concordant with its formation of a disulfide-bonded dimer.

Pre-titrated amounts of each sEC1-4 protein bound ANDV Gn/Gc; however, dimeric sEC1-4(T141C/G337C) appeared to recognize ANDV Gn/Gc less efficiently than its WT and obligate-monomer counterparts (**Figures 5G–H**). Together, these findings suggest that both monomeric and dimeric forms of PCDH1 provide suitable entry receptors for hantaviruses but also raise the possibility that viral particles may preferentially engage PCDH1 monomers.

### PCDH1 mutations disrupting Gn/Gc interaction protect Syrian hamsters against lethal ANDV challenge

We previously demonstrated that ANDV infection and virulence was highly attenuated in Syrian hamsters engineered to lack PCDH1 expression (6). Although these studies identified PCDH1 as a critical requirement in ANDV multiplication and pathogenesis *per se*, they did not specifically address if this requirement arose from PCDH1’s role as an entry receptor that directly engages the viral glycoprotein complex. To test the latter hypothesis, we used CRISPR/Cas9 genome engineering to introduce a F83A/D85R double mutation at the *PCDH1* locus in Syrian hamsters (**Figure 6A**). Biallelic *PCDH1(F83A/D85R)* animals expressed levels of PCDH1 in the lung similar to their WT counterparts (**Figure 6B**).

**Figure 6.**
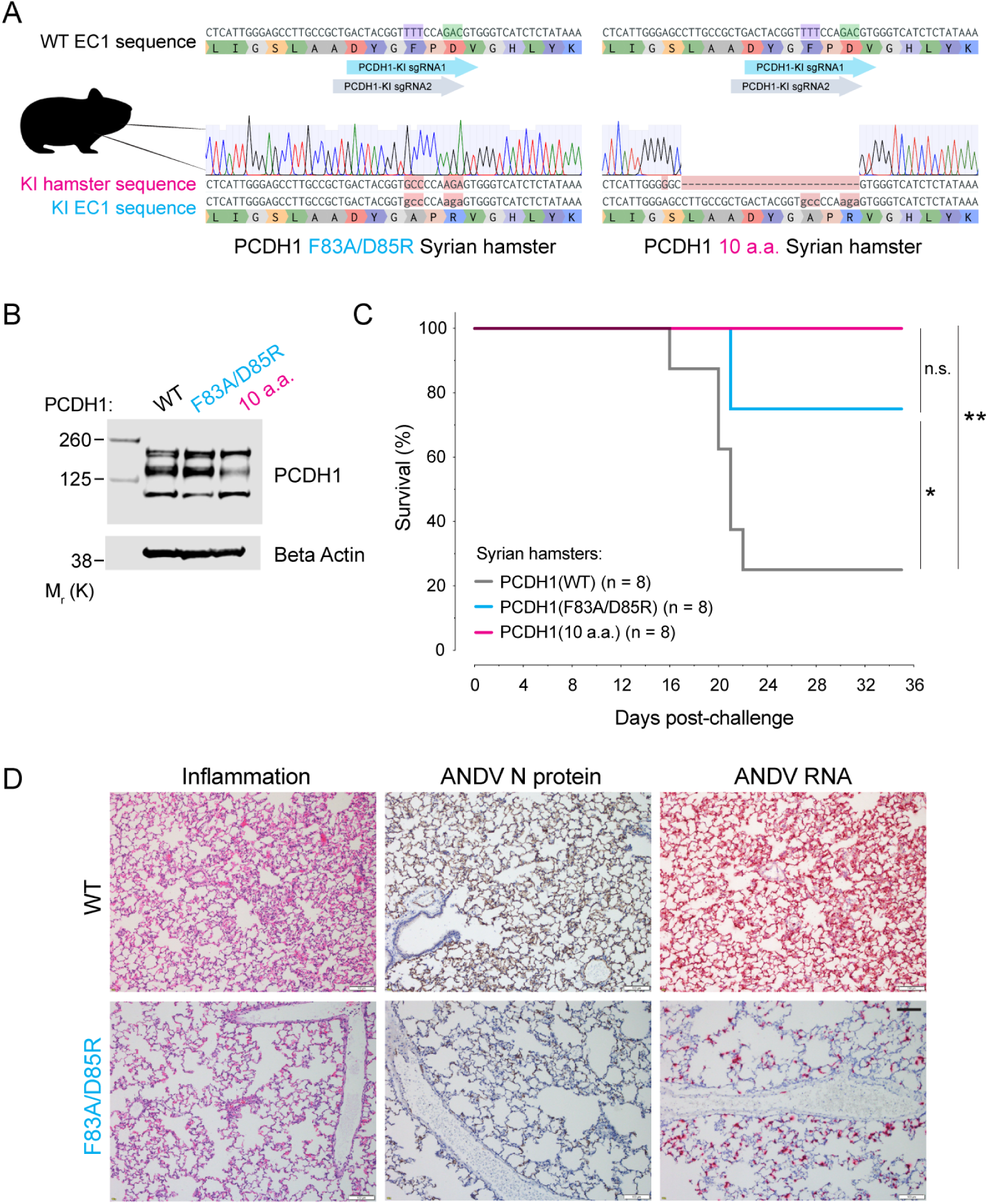
Two point mutations in PCDH1 confer protection of Syrian hamsters against a lethal ANDV challenge. (A) Reference nucleotide and amino acid sequence of PCDH1-EC1 Syrian hamster (WT, above) and representative sequences and trace files of Syrian hamsters after CRISPR-Cas9 genome editing [PCDH1(F83A/D85R), lower left] and [PCDH1(10a.a.) lower right]. The nucleotides encoding the corresponding human PCDH1-EC1 Gn/Gc-interacting residues, are highlighted: F83 in purple and D85 in green along with the location of the single guide RNAs (KI, knock-in; sgRNA, single guide RNA). (B) Immunoblot detecting PCDH1 in lung tissue lysates from WT or CRISPR knock-in mutant Syrian hamsters. Antibody targets PCDH1’s cytoplasmic tail. A representative blot of two independent experiments is shown. (C) Syrian hamster ANDV challenge. Groups of WT, PCDH1(F83A/D85R), and PCDH1(10a.a.) CRISPR knock-in mutant hamsters were inoculated intranasally with ANDV (2,000 PFU). Mortality was monitored and hamsters were euthanized on day 35 post-exposure. One experiment was performed, with *n* = 8 for each group. Data was analyzed using two-sided, log-rank Mantel-Cox test; *ns* > 0.05, \**P* < 0.05, \*\**P* < 0.01. (D) Lung sections from WT and PCDH1(F83A/D85R) hamsters were collected 15 days post ANDV exposure. Representative histochemical images indicate inflammation in pulmonary tissue (left), ANDV nucleoprotein (N) (middle, tan staining), and ANDV RNA (right, red staining, detected by in situ hybridization). Scale bars represent 100 µm.

Interestingly, we also identified founder animals with an allele encoding a larger 10 amino acid deletion in the Gn/Gc-binding surface: *PCDH1(S76G, ΔL77–D85)* (**Figure 6A**). This fortuitous mutant (hereafter, PCDH1(10a.a.)) abrogated PCDH1’s receptor function *in vitro*, but at the expense of a partial reduction in its steady state expression level, which was also observed *in vivo* (hamster lung tissue) (**Figures 6B and S7A, C**). Nevertheless, PCDH1(10a.a.)-expressing cell subpopulations, sorted to approximate WT PCDH1 expression levels, remained resistant to viral entry (**Figures S7B–C**) suggesting that the loss of 7 of 11 residues implicated in ANDV and SNV Gn/Gc recognition, and not reduced PCDH1 expression, accounts for PCDH1(10a.a.)’s loss of receptor activity. Considering that *PCDH1(10aa)* afforded a genotype intermediate to that of the *PCDH1(F83A/D85R)* and *PCDH1(KO)* alleles, we evaluated both CRISPR/Cas9 knock-in Syrian hamsters in ANDV challenge.

We challenged WT and knock-in hamsters intranasally with a lethal dose of ANDV and monitored their survival for up to 35 days post-exposure (**Figures 6C** **and** **S8**). Hamsters carrying modified *PCDH1* alleles (either *PCDH1(F83A/D85R)* or *PCDH1(10a.a.)*) were largely protected against a lethal outcome of ANDV challenge while the control animals carrying WT *PCDH1* succumbed to infection. Histochemical analysis of lung tissues at 15 days post ANDV exposure revealed reduced inflammation and expansion of alveolar septa due to monocyte infiltration (left panels), lower viral RNA (red staining, right panels) and nucleoprotein levels (tan staining, middle panels) in hamsters expressing PCDH1(F83A/D85R) compared to WT hamsters (**Figure 6D**). Seroconversion, indicative of an ANDV-specific IgG response, was observed in all surviving hamsters, indicating they had all been exposed to the virus (**Figure S8**). Our Syrian hamster challenge study strongly supports our hypothesis that a direct Gn/Gc:PCDH1 engagement is a critical requirement for ANDV multiplication and lethal, *in vivo* HCPS-like pathogenesis.

## Discussion

Virus-receptor interactions can influence viral entry, cell and tissue tropism, viral host range and pathogenesis, and afford attractive targets for antiviral countermeasures (22-31). We previously demonstrated that PCDH1 is an essential entry host factor for New World hantaviruses (6,7). Herein, we identified residues in the EC1 domain of PCDH1 critical for ANDV and SNV Gn/Gc engagement, including one that influences cellular host range, and demonstrated that Gn/Gc:PCDH1 recognition, not just PCDH1 *per se*, is required for development of lethal ANDV infection in Syrian hamsters. We conclude that PCDH1 is a *bona fide* entry receptor for ANDV and SNV, the primary etiologic agents of HCPS in the Americas, and is likely to play such a role for other New World hantaviruses shown to utilize PCDH1 for entry (6,32).

We leveraged the recently published crystal structure of PCDH1 domains 1-4 (EC1-4) (17) and structure-based prediction of interfacial residues to generate a list of candidate virus-contacting residues in PCDH1 EC1. Quantitative assessment of this panel of site-directed EC1 mutants, through receptor-binding and -blocking assays, showed that a membrane-distal EC1 surface centered around a flexible loop makes key contacts with Gn/Gc during hantavirus-receptor recognition (**Figures 2-3**). The identified key residues might either directly contact Gn/Gc or might indirectly contribute to Gn/Gc binding through intramolecular interactions that maintain the local geometry of EC1, as is likely the case for more buried residues (e.g., L152). Our findings also shed light on differences in the mechanisms of PCDH1 recognition by ANDV and SNV Gn/Gc. While ANDV Gn/Gc binds with higher avidity to PCDH1 than its SNV counterpart (**Figures 1H-I**) (6), both Gn/Gcs displayed similar binding patterns toward our PCDH1 mutant panel (**Figure 4A**), suggesting that they largely share their key PCDH1 contacts. Instead, the differences in binding affinity between ANDV and SNV Gn/Gcs (and by extension, the glycoproteins from other New World hantaviruses) likely arise from sequence variations in Gn/Gc’s yet-unmapped PCDH1-binding site (also see below). We further speculate that more divergent residues at those or adjacent positions with Gn/Gc account for the failure of Old World hantavirus Gn/Gcs to engage with PCDH1. Whether non-PCDH1–using Old World hantaviruses recognize their putative receptors through the same or distinct surfaces on Gn/Gc remains to be determined.

We provide herein the first evidence that hantavirus Gn/Gc:PCDH1 recognition can impact cellular host range. Specifically, we observed that murine endothelial cells are refractory to SNV Gn/Gc-dependent entry (**Figure 1A**) and mapped this human-murine difference in susceptibility to a sequence variation at a single residue in EC1, residue 83, that modulates Gn/Gc-PCDH1 engagement (**Figures 1H-I** **and S1B**). This PCDH1 ortholog-dependent effect on entry was not observed for ANDV, despite a discernible effect on receptor binding avidity (**Figures 1A, H-I**, **and S1B**), presumably because ANDV Gn/Gc, unlike its SNV counterpart, retains sufficient binding avidity for murine PCDH1 on cell surfaces. Residue F83 afforded us a nidus to more comprehensively map the Gn/Gc-interacting surface in EC1, and to identify 10 additional surface-exposed residues that are key for virus-receptor recognition. Indeed, mutation of F83 in combination with the adjacent D85 could further ablate PCDH1-dependent, ANDV Gn/Gc-mediated entry and infection *in vitro* (**Figure 4A**), and significantly reduced mortality from lethal ANDV challenge in Syrian hamsters (**Figure 6C**). Hamsters carrying a larger 10-amino acid deletion in PCDH1, encompassing F83, D85, and additional interfacial residues that individually and collectively impacted Gn/Gc-PCDH1 binding, phenocopied *PCDH1-*KO hamsters in being fully protected against lethal virus challenge (**Figure 6C**), likely by further reducing receptor recognition and cellular susceptibility to viral infection *in vivo*, as observed in cell culture (**Figure S7C**).

Our findings thus point to a role for New World hantavirus Gn/Gc:PCDH1 recognition in influencing viral host range in nature and suggest at least three new avenues for exploration, including: (i) sequencing and functional analyses of Gn/Gc (from circulating viruses) and *PCDH1* orthologs in geographically/ecologically distinct populations of rodents established as orthohantavirus hosts (e.g., the pygmy rice rat for ANDV); (ii) investigations of virus-receptor compatibility in natural hantavirus infections of rodents not traditionally considered to be viral reservoirs (e.g., SNV in wild-caught house mice (33)); and (iii) experimental cross-species infections to uncover novel virus-receptor mismatches that may uncover additional determinants of viral host range (34). Moreover, the virus-receptor interactions defined herein may illuminate our understanding of orthohantavirus infection and pathogenesis in the available animal models (35-41), and facilitate the development of new models to study virus-host interactions and test countermeasures. As a case in point, the New World hantaviruses ANDV and SNV do not appear to infect laboratory mouse strains, possibly due in part to the incompatibility between Gn/Gc and murine PCDH1 (especially for SNV) that we uncovered in this study. We speculate that transgenic mice bearing compatible *PCDH1(L83F)* alleles may sustain viral replication by overcoming a key host barrier to viral entry.

The disruption of PCDH1’s cellular function has been linked to the dysfunction of the airway epithelial barrier in respiratory diseases such as asthma (10). This raises the tantalizing possibility that the cellular function(s) of PCDH1 are intertwined with the pathogenesis of severe pulmonary disease caused by ANDV and SNV infections in humans. Although PCDH1’s endogenous functions and their molecular mechanisms remain poorly understood, its capacity to form *trans*-dimers through EC1:EC4 domain interactions is proposed to be central to its adhesive capacity (17). Here, we found that hantaviruses recognize a surface in PCDH1 EC1 that is distinct from the EC1-EC4 adhesive interface (**Figures 5A-B**), suggesting that virus-receptor interaction during entry, or putative glycoprotein-receptor interactions in infected cells, are unlikely to interfere directly with PCDH1 dimerization or vice versa. Studies with recombinant PCDH1 mutants engineered to form only monomers or only dimers supported this hypothesis (**Figure 6H**), as did the observation that deletion of EC4 (and abrogation of EC1-EC4 association) did not impair viral entry (**Figure 6C**). However, we did also obtain preliminary evidence that PCDH1 dimerization (or attendant conformational changes) may subtly disfavor Gn/Gc interaction (**Figure 6C**), raising the possibility that Gn/Gc expression in infected cells may shift the PCDH1 monomer-dimer equilibrium by preferentially sequestering monomers at the cell surface. More work is needed to test this idea as well as simpler (e.g., PCDH1 down-regulation) and more complex scenarios (e.g., Gn/Gc-induced changes in PCDH1 conformation or transmembrane signaling), through which hantavirus infection could perturb PCDH1’s endogenous functions in the mammalian airway.

## Supporting information

Supplemental Item 1

## Acknowledgments

We would like to thank Estefania Valencia, Laura Polanco and Isabel Gutierrez for technical support, Eva Mittler for valuable input, and the Einstein Flow Cytometry Core. Figures 1G and 3A were created with BioRender.com. This work was supported by NIH grant AI132633 (to K.C., J.M.D, and Z.W.), an Irma T. Hirschl/Monique Weil-Caulier Research Award, and by a Harold and Muriel Block Faculty Scholarship at the Albert Einstein College of Medicine (to K.C.).

## Author contributions

Conceptualization, M.M.S., A.S.H., Z.W., R.K.J., and K.C.; Methodology, M.M.S., R.L., A.S.H., R.K.J.; Software, G.L.; Validation, M.M.S., R.L., A.S.H., A.I.K., R.R.B., S.R.M., Y.L., A.G., A.M.M., X.Z., R.K.J.; Formal analysis, M.M.S., A.S.H., G.L.; Investigation, M.M.S., R.L., A.S.H., A.I.K., R.R.B., S.R.M., Y.L., A.G., A.M.M., X.Z., R.K.J.; Resources, R.L., Y.L.; Data curation, M.M.S., R.L., A.S.H., G.L., A.G., R.K.J.; Writing-original draft, M.M.S. and K.C.; Writing-review and editing, M.M.S., R.L., A.S.H., G.L., S.R.M., A.G., S.A., R.K.J., K.C.; Visualization, M.M.S., R.L., G.L., R.K.J., K.C.; Supervision, A.S.H., J.M.D., Z.W., K.C.; Project Administrator, K.C.; Funding Acquisition, K.C.

## Materials and methods

### Cells

Human osteosarcoma U2OS cells (ATCC) and *PCDH1*-knockout (KO) U2OS cells, generated as described in (6), were cultured in modified McCoy’s 5A media (Thermo Fisher), supplemented with 10% fetal bovine serum (FBS, Atlanta Biologicals), 1% GlutaMAX (Thermo Fisher), and 1% penicillin-streptomycin (Pen-Strep, Thermo Fisher). Human umbilical vein endothelial cells (HUVEC, Lonza) were cultured in EGM media supplemented with EGM-SingleQuots (Lonza). Human pulmonary microvascular endothelial cells (HPMEC, Promocell) and mouse primary lung microvascular endothelial cells (MLMEC, Cell Biologics) were cultured in MV2 Endothelial Cell Growth medium (Promocell) and Complete Mouse Endothelial Cell Medium (Cell Biologics), respectively. All adherent cell lines were maintained in a humidified 37°C, 5% CO_2_ incubator. Freestyle^TM^-293-F suspension cells (Thermo Fisher) were maintained in FreeStyle™ 293 expression medium (Thermo Fisher) using shaker flasks at 115 rpm, 37°C, and 8% CO_2_.

### rVSVs and infections

Replication-competent, recombinant vesicular stomatitis Indiana viruses (rVSVs) expressing an eGFP reporter and bearing hantavirus Gn/Gc of either ANDV (NP_604472.1), SNV (NP_941974.1), or HTNV (NP_941978.1) with the modifications described in (42), were generated using a plasmid-based rescue system in 293T cells and propagated on Vero cells as described previously (43,44). Viral infection was measured by automated enumeration of eGFP-expressing cells from captured images, using a custom analysis pipeline in CellProfiler (45), or directly from multiwell plates, using a CellInsight CX5 automated fluorescence microscope and onboard HCS Studio software (ThermoFisher) or Cytation 5 cell imaging multi-mode reader (BioTek). For sEC1-2 infectivity-inhibition experiments, pre-titrated amounts of rVSV particles were incubated with increasing concentrations of wild-type (WT) or mutant sEC1-2 at room temperature (RT) for 1 hour prior to addition to cell monolayers in 96-well plates. Number of infected cells was measured as above.

### Authentic hantaviruses and infections

ANDV strain Chile-9717869, SNV strain CC107, and HTNV strain 76-118 were propagated in Vero E6 cells as described previously (35,46). Hantavirus infections were performed, and infected cells were immunostained for viral antigen, as described previously (43). Briefly, U2OS *PCDH1*-KO cells, complemented with WT or mutant PCDH1, were exposed to virus at a multiplicity of infection (MOI) of 0.5-3 plaque forming units (PFU) per cell, and viral infectivity was determined by immunostaining of formalin-fixed cells at 72 h post-infection. Images were acquired at 20 fields per well, with a 20x objective on an Operetta high-content imaging device (PerkinElmer). Images were analyzed with a customized scheme built from image analysis functions present in Harmony software and the percentage of infected cells was determined using the analysis functions.

### PCDH1 EC1 loop modeling

The EC1 missing loop (residues 80-89) in the crystal structure of human PCDH1 was modeled using MODELLER through the Chimera plugin (17,47,48). All 10 modeled loop conformations scored similarly, based on their GA341 and zDOPE scores (49,50). The two most divergent loop conformations were selected as representative loop conformations to perform structure-based interfacial residue predictions on.

### Interfacial residue predictions in the PCDH1 ectodomain

Five complimentary structure-based tools (51-55) were used to predict interfacial residues in the PCDH1 ectodomain crystal structure, with two modeled alternative conformations for the missing loop (residues 80-89; see above). For any given prediction method, the results for each of the two versions of the PCDH1 ectodomain structure (Pred_A_, Pred_B_) were combined into a single set of predictions (Pred_A_ ∪ Pred_B_). Predicted interfacial residues were then ranked based on the number of supporting algorithms (range 0-5).

### Cloning soluble PCDH1 variants

Constructs encoding soluble (secreted) PCDH1 variants (sEC1-2 and sEC1-4) were generated by cloning the following sequences into the pcDNA3.1 mammalian expression vector (ThermoFisher): EC1-EC2 (residues 1-284) (6) or EC1-EC4 (residues 1-503), each in frame with a C-terminal GSG linker, followed by Myc, Flag, and deca-histidine tags. Each construct also retained the endogenous PCDH1 N-terminal signal sequence (residues 1-57). PCDH1 point mutations were cloned into the pcDNA3.1 plasmid using standard molecular biology techniques. To avoid free cysteine-mediated, non-specific protein-protein interactions, sEC1-4 variants included a C432S mutation. The C432S mutation did not have an observable effect on the production of the soluble protein or on the binding to rVSV-ANDV-Gn/Gc. The sequences of all the plasmid inserts were confirmed by Sanger sequencing.

### Expression and purification of soluble PCDH1 variants

Soluble PCDH1 variants cloned into pcDNA3.1 (see above) were expressed in 293F cells in shaker flasks by transient transfection with linear polyethyleneimine (Polysciences), as described previously (56), and purified by nickel-chelation chromatography. Cell supernatants were clarified and stirred overnight at 4°C with proteinase inhibitor (Sigma-Aldrich) and nickel-NTA resin (Qiagen) at 0.3 mL packed resin per 50 mL cell supernatant. Nickel-NTA beads were then collected, washed with phosphate buffer saline (PBS) containing 50 mM imidazole, and eluted with PBS containing 250 mM imidazole. The eluted protein was buffer-exchanged with PBS, concentrated, and stored in aliquots at 80°C. The purity of the secreted PCDH1 variants was determined by size-exclusion chromatography (SEC) and/or either SDS-PAGE or Native-PAGE gels, stained with Bio-Safe™ Coomassie G-250 Stain (Bio-Rad) and imaged on a Li-Cor Odyssey^Ⓡ^ Fc Imager (Li-Cor Biosciences). For analytical SEC, a Superdex S200 (10/300) column was equilibrated in PBS and calibrated with Gel Filtration Standard (Bio-Rad) composed of thyroglobulin (MW 670 kDa), bovine γ-globulin (MW 158 kDa), chicken ovalbumin (MW 44 kDa), horse myoglobin (MW 17 kDa), and vitamin B12 (MW 1.35 kDa).

### Monoclonal antibody:PCDH1 binding ELISA

To determine the capacity of an infection-inhibiting, EC1-specific monoclonal antibody (mAb) to bind to sEC1-2 variants, ELISA plates were coated with serial twofold dilutions of WT or mutant sEC1-2 (starting concentration at 400 ng/well) overnight at 4°C. After briefly washing with PBS, wells were blocked with 5% nonfat dry milk in PBS (1 h at RT) and incubated with anti-EC1 mAb-3305 (1 µg/mL, 1 h at RT). After washing with PBS, bound antibody was detected by incubation with an anti-human HRP antibody (Thermo Fisher), 0.045 µg/mL, 1 h at RT). ELISA signal was developed using 1-Step^TM^ Ultra TMB-ELISA substrate solution (Thermo Scientific) and measured at an absorbance of 450 nm on a Perkin Elmer Wallac 1420 Victor2^TM^ microplate reader or Cytation 5 cell imaging multi-mode reader (BioTek).

### rVSV:PCDH1 competition ELISA

The capacity of sEC1-2 mutants to compete with the binding of sEC1-2(WT) to rVSVs, bearing ANDV or SNV Gn/Gc, was determined by a competition capture ELISA. High-protein binding 96-well ELISA plates (Corning) were coated with purified sEC1-2 (100 ng /well) overnight at 4°C, washed briefly with PBS, and blocked with 5% nonfat dry milk in PBS (1 h at RT). Pre-titrated amounts of rVSV-ANDV-Gn/Gc or rVSV-SNV-Gn/Gc, were membrane-labeled with a short-chain phospholipid probe, functional-component spacer diacyl lipid conjugated to biotin (FSL-biotin; Sigma-Aldrich), as described previously (43). The rVSVs were pre-incubated with serial 3x-diluted sEC1-2(WT or variants) for 1 h at RT prior to their incubation with sEC1-2 coated wells (1 h at 37°C). Bound rVSVs were detected by incubation with a streptavidin-horseradish peroxidase (HRP) conjugate (Thermo Scientific). ELISA signal was developed using 1-Step^TM^ Ultra TMB-ELISA substrate solution (Thermo Scientific) and measured at an absorbance at 450 nm on a Perkin Elmer Wallac 1420 Victor2^TM^ microplate reader or Cytation 5 cell imaging multi-mode reader (BioTek).

### rVSV:PCDH1 binding ELISA

The capacity of rVSVs bearing ANDV Gn/Gc to recognize WT, monomer, and dimer sEC1-4 was determined by capture ELISA. High-protein binding 96-well ELISA plates (Corning) were coated with purified sEC1-4 (100 ng /well) overnight at 4°C, washed briefly with PBS, and blocked with 5% nonfat dry milk in PBS (1 h at RT). Pre-titrated amounts of rVSV-ANDV-Gn/Gc were membrane-labeled with FSL-biotin as described previously (43), and serially diluted (twofold). rVSVs were added to sEC1-4 coated plates and incubated for 1 h at 37°C. Bound rVSVs were detected by incubation with a streptavidin-HRP conjugate (Peirce^TM^, and sEC1-4 protein was detected by incubation with an anti-Flag m2 mAb-HRP conjugate (Sigma-Aldrich) (1 h at 37°C). ELISA signal was developed and measured as noted above.

### Hierarchical data clustering

An in-house algorithm was used to perform hierarchical clustering of two experimental readouts: sEC1-2 competition ELISA (absorbance at 450 nm) and infection-inhibition assay (GFP positive cells [%]). A sigmoidal function [1] (see below) was fitted to the normalized experimental readouts using a non-linear least square analysis as implemented in the SciPy package (57). Hierarchical clustering and a heatmap comparing each sigmoidal curve to all other sigmoidal curves were built using the clustermap (method = average; metric = Euclidean distance) function as implemented in the seaborn python package (https://seaborn.pydata.org/index.html).

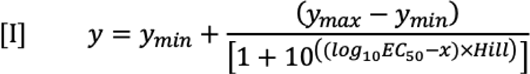

Where y corresponds to the experimental readout (ER) (relative ELISA signal or percent infectivity); y_min_ and y_max_ are the minimum and maximum ERs, respectively; EC_50_ is the value that gives half-maximum ER (half-y_max_); Hill describes the slope of the curve, and x is the amount of soluble protein, log_10_(sEC1-2), at the particular y (ER). (Original code can be found at: https://github.com/chandranlab/pcdh1_interface.git).

### Generation of stable cell populations expressing PCDH1 variants

cDNA constructs encoding full-length human PCDH1 (isoform 1, Genbank accession number NM_002587) or lacking the fourth extracellular cadherin (EC) repeat ΔEC4 (residues 328-446) were synthesized in frame with Myc and Flag epitope tags at the C-terminus (Epoch Biolabs or Twist Bioscience) and cloned into the pBABE-puro retroviral vector (58). Residue numbers are based on the full-length sequence of PCDH1 starting from the signal sequence (i.e., residue D85 in this study matches pdb 6MGA residue D28, and E137 in this study matches pdb 6MGA residue E80). Mutations for the PCDH1 variants were introduced into the pBABE-puro-PCDH1 plasmid using standard molecular techniques and confirmed by Sanger sequencing. Human U2OS *PCDH1*-KO cells ectopically expressing the above PCDH1 variants were generated by transduction with pBABE-puro-based retroviral vectors. Retroviruses packaging the transgenes were produced by transfecting 293T cells (43), and target cells were directly exposed to sterile-filtered, retrovirus-laden supernatants in the presence of polybrene (6 μg/mL). Transduced U2OS cell populations were selected with puromycin (2 μg/mL), and transgene expression was confirmed by immunostaining. HPMEC and mouse pulmonary endothelial cells were also transduced as above but were not subjected to antibiotic selection.

### Detection of PCDH1 surface expression by flow cytometry

Human U2OS cells expressing variant PCDH1 were seeded in 6-well plates 24 hours prior to immunostaining. Cells were chilled on ice for 10 min and blocked with chilled PBS/10% FBS for 30 min at 4°C. Surface PCDH1 was stained using human anti-EC7 mAb-3677 (5 µg/mL) followed by anti-human Alexa Fluor^TM^ 555 (ThermoFisher) for 1 h at 4°C each. After washing, cells were stained with Live/Dead^TM^ Fixable Violet Dead Cell Stain Kit (Invitrogen), washed with PBS, and re-suspended in PBS/2% FBS. Stained cells were passed through a 0.41 µm Nylon Net Filter (Millipore) and analyzed using an LSRII Flow Cytometer (BD Biosciences) and FloJo V.10 software. Subpopulations of cells expressing PCDH1(10a.a.) were isolated by FACS (WOLF Cell Sorter, NanoCellect) and verified following the staining method described above.

### Immunofluorescence microscopy of PCDH1 surface expression

Human U2OS cells expressing variant PCDH1 were seeded on fibronectin-coated glass coverslips 24 h pre-immunostaining. Cells were washed briefly in PBS before blocking with chilled PBS/10% FBS for 30 min at 4°C. PCDH1 was detected by a human anti-EC7 mAb-3677 (5 µg/mL) or human anti-EC1 mAb-3305 (5 µg/mL) for 1 h at 4°C. After washing with chilled PBS, the cells were fixed with 4% formaldehyde (Sigma-Aldrich) for 5 min followed by staining with an anti-human Alexa Fluor^TM^ 555 antibody (ThermoFisher). Coverslips were mounted on glass slides using ProLong^TM^ Gold Antifade Mountant containing DAPI (ThermoFisher) and cells were examined using a Axio Observer Z1 wide field epifluorescence microscope (Zeiss Inc.) with a 40x objective. Images were processed in Photoshop software (Adobe Systems).

### Animal welfare statement

Breeding, CRISPR/Cas9 genome engineering, and challenge studies with Syrian hamsters were conducted under IACUC-approved protocols in compliance with the Animal Welfare Act, PHS Policy, and other applicable federal statutes and regulations related to animals and experiments involving animals. The facilities where this research was conducted (Utah State University and USAMRIID) are accredited by the Association for Assessment and Accreditation of Laboratory Animal Care, International (AAALAC), and adhere to principles stated in the Guide for the Care and Use of Laboratory Animals, National Research Council, 2011.

### Generation of *PCDH1*-gene edited Syrian hamsters by CRISPR/Cas9 genome engineering

A panel of candidate sgRNAs was designed, assembled by overlapping PCR to generate human U6 promoter-driven sgRNA expression cassettes, and screened for genome-editing efficiency in BHK21 baby hamster kidney cells stably expressing Cas9. The best candidate sgRNA (sgRNA2: 5’-GACTACGGTTTTCCAGACTGGG-3’) targeted sequences encoding the homologs of human F83 and D85 in the hamster *PCDH1* gene (F79 and D81, respectively) (accession number NW_024429184.1). In addition, a knock-in, single donor strand (sequence: 5’-CCAACACCCTCATTGGGAGCCTTGCCGCTGAC TACG<u>GT**gcc**C</u>CA**aga**GTGGGTCATCTCTATAAACTAGAGGTAGGTGCTCCATATCTTC-3’ with the BaeGI cleavage site underlined and the bold, lowercase letters indicating the F83A and D85R mutation sites) was used for *in vivo* gene editing. The sgRNA was *in vitro*-transcribed and assembled into sgRNA/Cas9 ribonucleoprotein complexes, diluted with 10 mM RNase-free TE buffer to a concentration of 50 ng/μL sgRNA, 5uM single donor strand, and 50 ng/μL Cas9, for pronuclear injections. PCDH1 gene edited hamsters were produced following the procedure described in (6). Genomic DNA was isolated from hamster pups at the age of 2 weeks, a product flanking the sgRNA target sites was PCR-amplified and subjected to a T7 Endonuclease I assay (NEB) to detect indels. Amplicons from pups bearing indels were TOPO-cloned and sequenced to identify founder animals carrying nucleotide changes corresponding to the F83A and D85R mutations, in addition to the “10 amino acid” labeled founder animal, with a S76G mutation and a deletion of residues L77–D85 (corresponding to the homologous hamster residues S72 and L73-D81, respectively).

### Western blot on hamster lung tissues

PCDH1 expression in *PCDH1*-KI Syrian hamsters (described above) was confirmed by immunoblotting lung homogenates from WT and *PCDH1*-KI hamsters. Hamster lungs were placed in Cell Lysis Buffer (Invitrogen), containing 1 mM PMSF and protease inhibitor (Sigma-Aldrich), before homogenizing with zirconium beads. The supernatant was collected and the total amount of protein was determined via Bradford assay (BioRad). 30 µg of protein was added and verified using a mouse anti-BetaActin mAb (CellSignaling). PCDH1 was detected using a polyclonal antibody targeting the PCDH1 cytoplasmic tail (ThermoFisher).

### Syrian hamster challenge studies

Groups of wild-type Syrian golden hamsters (Envigo) and *PCDH1-*KI (*PCDH1(10a.a.)* or *PCDH1(F83A/D85R*)) Syrian golden hamsters (Utah State University), 5-12 weeks old, male and female, were exposed to 2,000 PFU of ANDV strain Chile-9717869 diluted in PBS, via the intranasal route and monitored for up to 35 days post-exposure. Animals were observed daily for clinical signs of disease, morbidity, and mortality. Moribund animals, described as being unresponsive or presenting with severe respiratory disease, were humanely euthanized on the basis of IACUC-approved criteria.

### Histopathology, immunohistochemistry, and in situ hybridization

Lungs harvested from *PCDH1(WT)* and *PCDH1(F83A/D85R)* hamsters (*n* = 3), 15 days post ANDV exposure, were fixed in buffered formalin for 30 days. Lung tissues were removed from biocontainment and processed at the USAMRIID histology lab. The tissues were trimmed, processed, embedded in paraffin, cut by microtomy, stained, coverslipped, and screened. For histopathology, 5-µm-thin sections were cut and stained with haematoxylin and eosin using standard procedures.

Immunohistochemistry was performed using the Dako Envision system (Dako Agilent Pathology Solutions). Briefly, after deparaffinization, peroxidase blocking, and antigen retrieval, sections were covered with a rabbit polyclonal anti-Sin Nombre virus antibody (#1244, USAMRIID) that cross binds to Andes virus at a dilution of 1:5,000 and incubated at RT for 40 minutes (mins). Sections were then rinsed, and a peroxidase-labeled polymer (secondary antibody) was applied for 40 mins. Slides were rinsed and a brown chromogenic substrate 3,3’ Diaminobenzidine (DAB) solution (Dako Agilent Pathology Solutions) was applied for eight mins. After the substrate-chromogen solution was rinsed off, the slides were counterstained with hematoxylin and rinsed. The sections were dehydrated, cleared with Xyless, and then coverslipped.

In-situ hybridization (ISH) was performed to detect ANDV RNA using the RNAscope 2.5 HD RED kit (Advanced Cell Diagnostics) according to the manufacturer’s instructions. An ISH probe targeting ANDV S segment (genbank accession number: NC_003466.1) was designed and synthesized by Advanced Cell Diagnostics (#900241, Advanced Cell Diagnostics). Tissue sections were deparaffinized with xylene, underwent a series of ethanol washes and peroxidase blocking, and heated in kit-provided, antigen retrieval buffer followed by digestion by kit-provided protease. Sections were exposed to ISH target probe pairs and incubated at 40°C in a hybridization oven for 2 h. After rinsing, the ISH signal was amplified using a kit-provided Pre-amplifier and Amplifier conjugated to alkaline phosphatase and incubated with a Fast Red substrate solution for 10 mins at RT. Sections were then counterstained with hematoxylin, air-dried, and coverslipped.

### Serology of hamsters following ANDV challenge

ANDV Gn/Gc-specific IgG titers were determined by end-titer ELISA using rVSV-ANDV-Gn/Gc. Briefly, high-protein binding 96-well ELISA plates (Corning) were coated with 10 μg/ml rVSV-ANDV-Gn/Gc, diluted in PBS, and incubated overnight at 4°C. Wells were then blocked in 5% milk protein in PBS/0.02% Tween 20 (2 h at RT). Serum samples were serially diluted in 5% milk protein in PBS/0.02% Tween 20 and added to antigen-coated plates (2 h at RT). Plates were washed with PBS/0.02% Tween20 before adding HRP-conjugated goat anti-hamster IgG (Seracare Life Sciences) (1 h at RT). Following a final wash, 2,2’-Azinobis [3-ethylbenzothiazoline-6-sulfonic acid]-diammonium salt (ABTS) substrate (Kirkegaard and Perry Laboratories, Inc.) was added and absorbance values were read at 405 nm using a Spectramax® plate reader (Molecular Devices, LLC).

---

**Figure S1.**
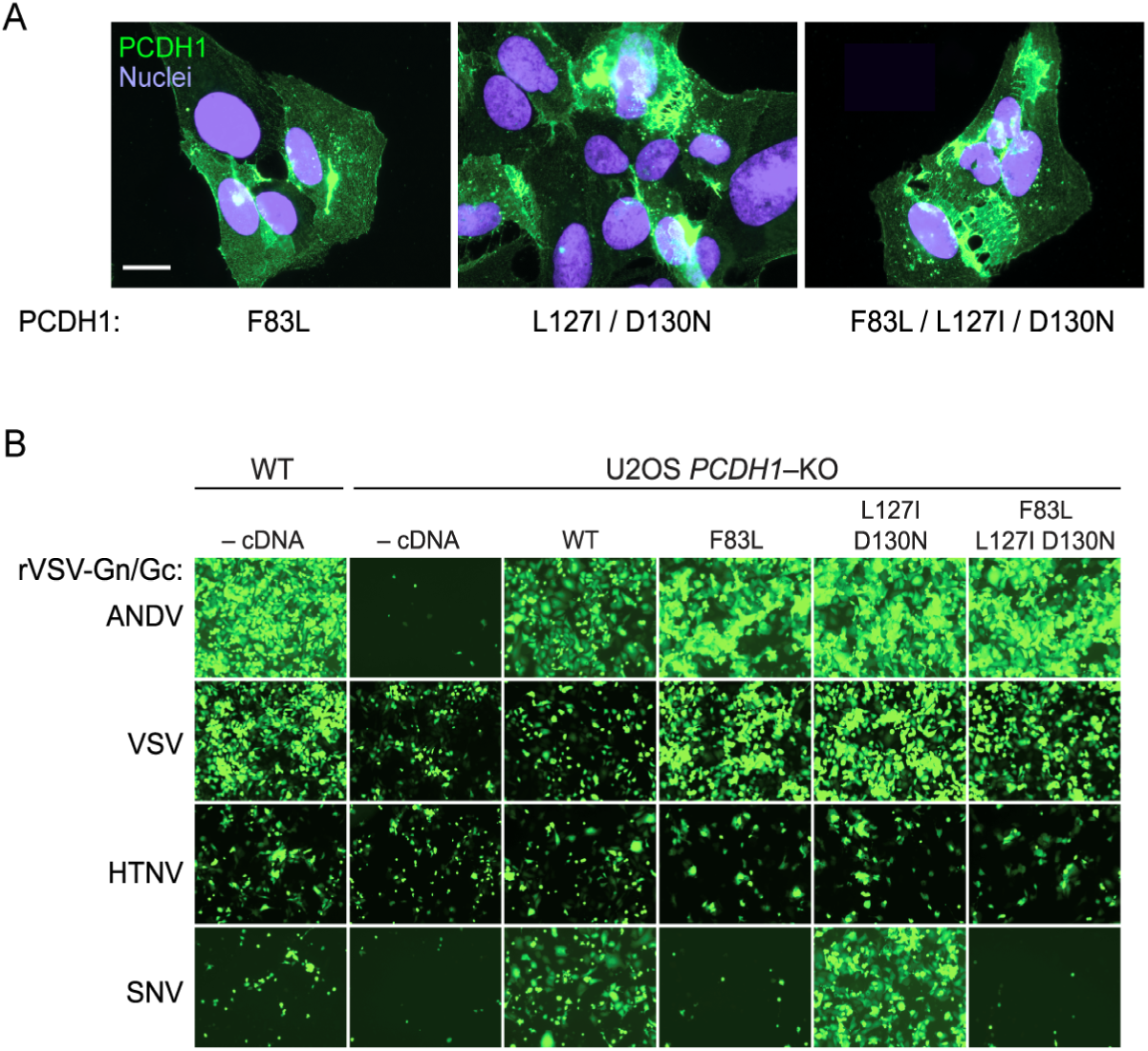
Residue F83 in PCDH1 is a key determinant of Sin Nombre virus infection. (A) Expression of PCDH1 variants. U2OS *PCDH1*-KO cells expressing the indicated PCDH1 variants were immunostained with an anti-flag antibody. Scale bar, 20 µm. (B) Capacity of human PCDH1 cell variants bearing EC1 mutations to support hantavirus Gn/Gc dependent entry. U2OS *PCDH1*-KO cells expressing the indicated PCDH1 variants were exposed to rVSVs bearing the indicated Gn/Gc proteins. -cDNA, no complementing PCDH1. eGFP-positive cells (pseudocolor green) were detected by immunofluorescence microscopy.

**Figure S2.**
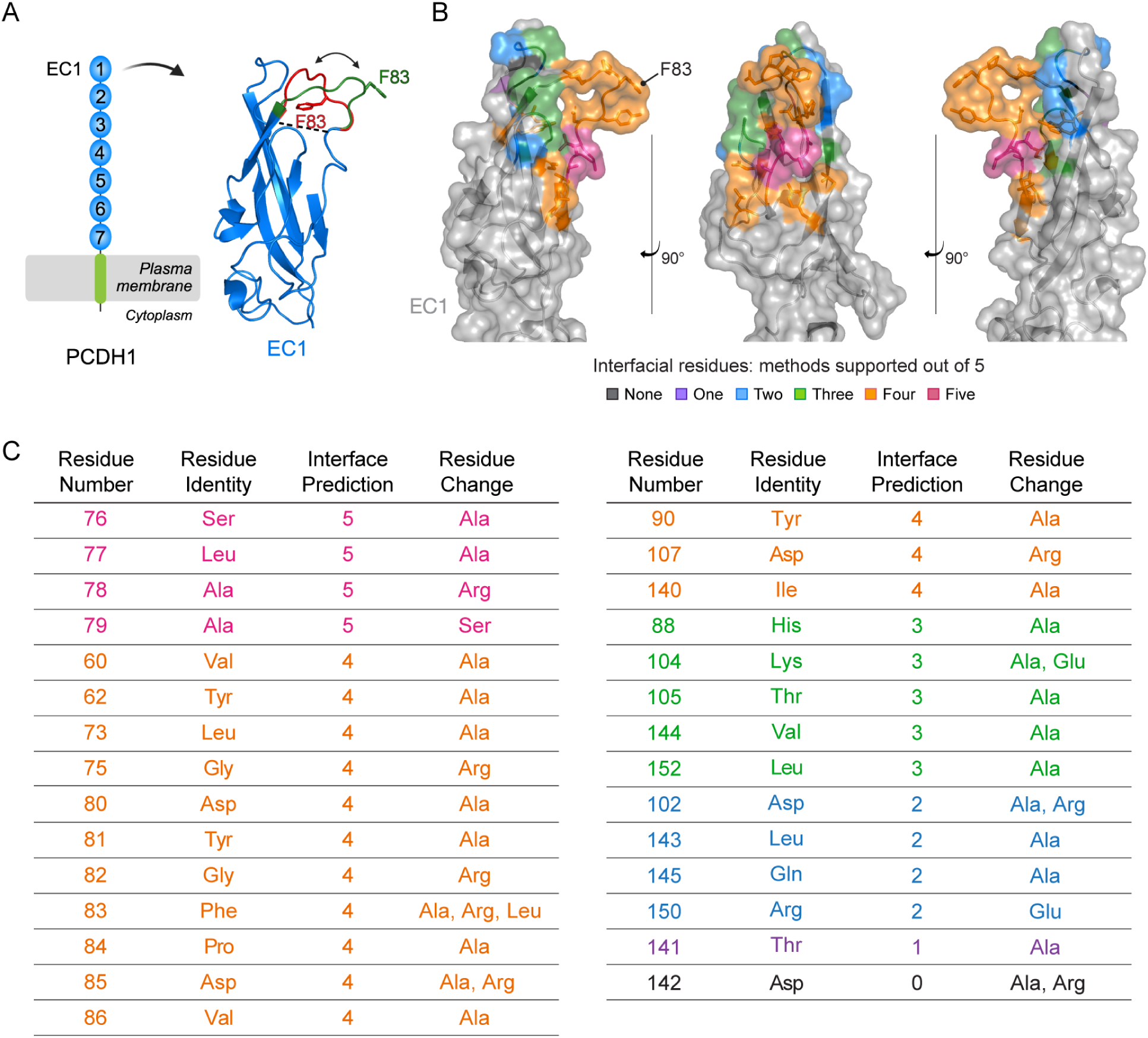
Structure-based interfacial prediction reveals a surface patch on PCDH1 EC1 that potentially drives the interaction with ANDV and SNV Gn/Gc. (A) Schematic representation of PCDH1 and crystal structure of EC1 (PDB 6MGA) displaying two modeled conformations (green: “open conformation”, red: “closed conformation”) for the disordered, uncrystallised loop comprising of residues 80-89. Residue F83 is indicated in each predicted loop conformation. (B) EC1 crystal structure in the “open conformation” displaying the EC1 residues chosen for mutational screening, ranked according to the number of supporting algorithms. Structure adapted from PDB 6MGA. (C) List of the EC1 residues chosen for mutational screening in (B) ranked according to the number of supporting algorithms (interface prediction column). The amino acid substitution(s) are listed for each residue.

**Figure S3.**
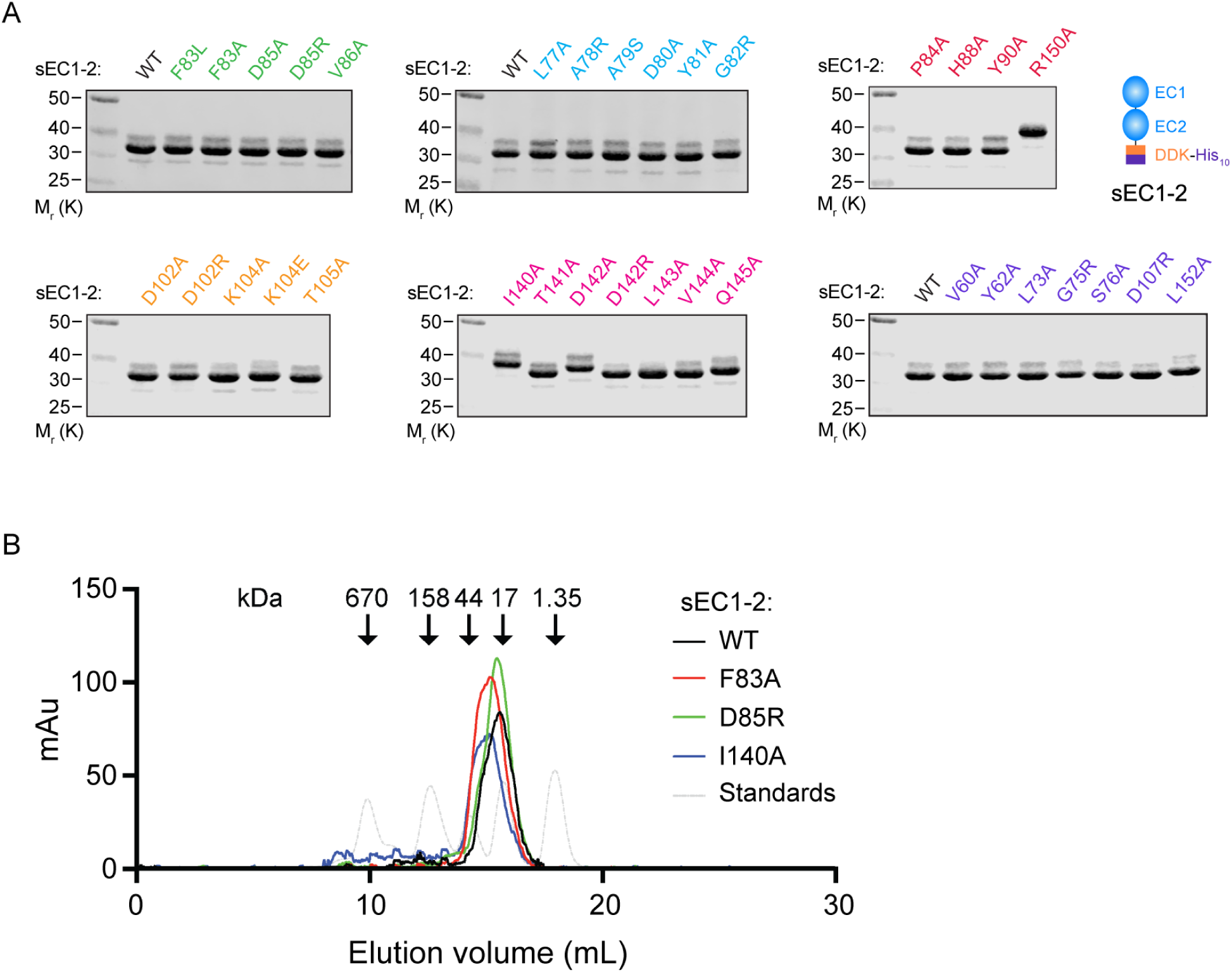
Expression and purification of PCDH1 soluble EC1-2 mutants. (A) Purified WT and mutant sEC1-2 proteins were separated on an SDS-polyacrylamide gel and visualized by Coomassie Brilliant Blue staining. M_r_, relative molecular weight (K denotes x 1,000). Schematic representation of sEC1-2 (right). (B) The elution profile of size exclusion chromatography (SEC) of WT and a selection of mutant sEC1-2 proteins from (A). Absorbance (milli arbitrary units, mAu) was measured at 280 nm using a calibrated Superdex S200 column at a physiological salt concentration. The dotted gray line shows elution profiles of SEC standards (molecular weight indicated) under the same buffer conditions.

**Figure S4.**
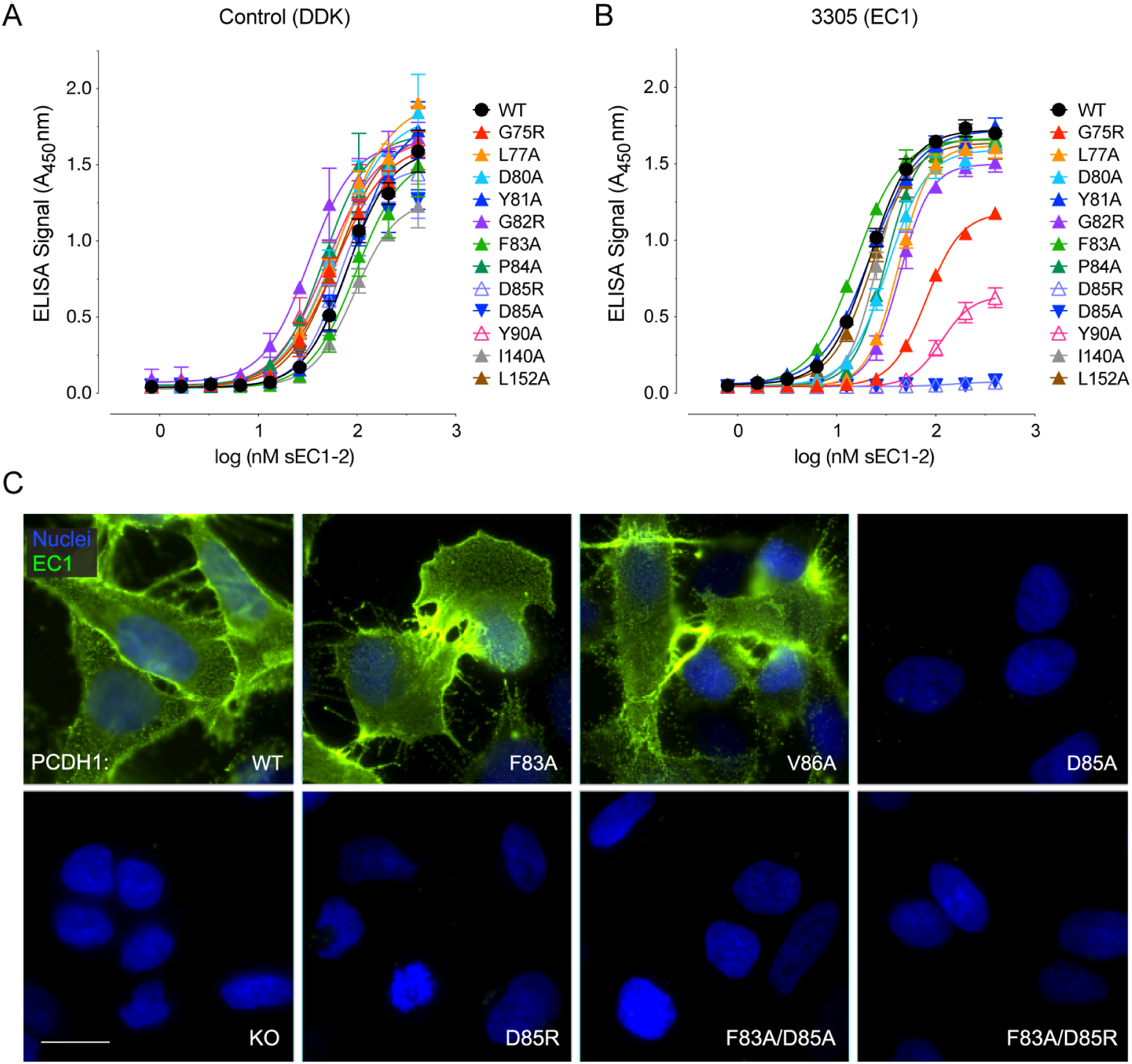
New World hantavirus infection-blocking 3305-mAb recognizes PCDH1 EC1 residue D85. (A) ELISA detecting Flag-tagged WT and mutant sEC1-2 proteins, coated on plates, using an anti-Flag-HRP antibody. Averages ± SD: two experiments, *n* = 3-4. (B) The capacity of anti-EC1 3305-mAb to bind to WT or mutant sEC1-2 proteins. Averages ± SD: two experiments, *n* = 4. Done in parallel with (A). (C) Detection of PCDH1 variants using anti-EC1 3305-mAb on U2OS *PCDH1*-KO cells complemented with WT or mutant PCDH1. Although the cells expressing PCDH1(D85) variants are not stained here, the protein is expressed in these cells (Figure S5) using an EC7-specific, 3677-mAb. Scale bar, 20 µm.

**Figure S5.**
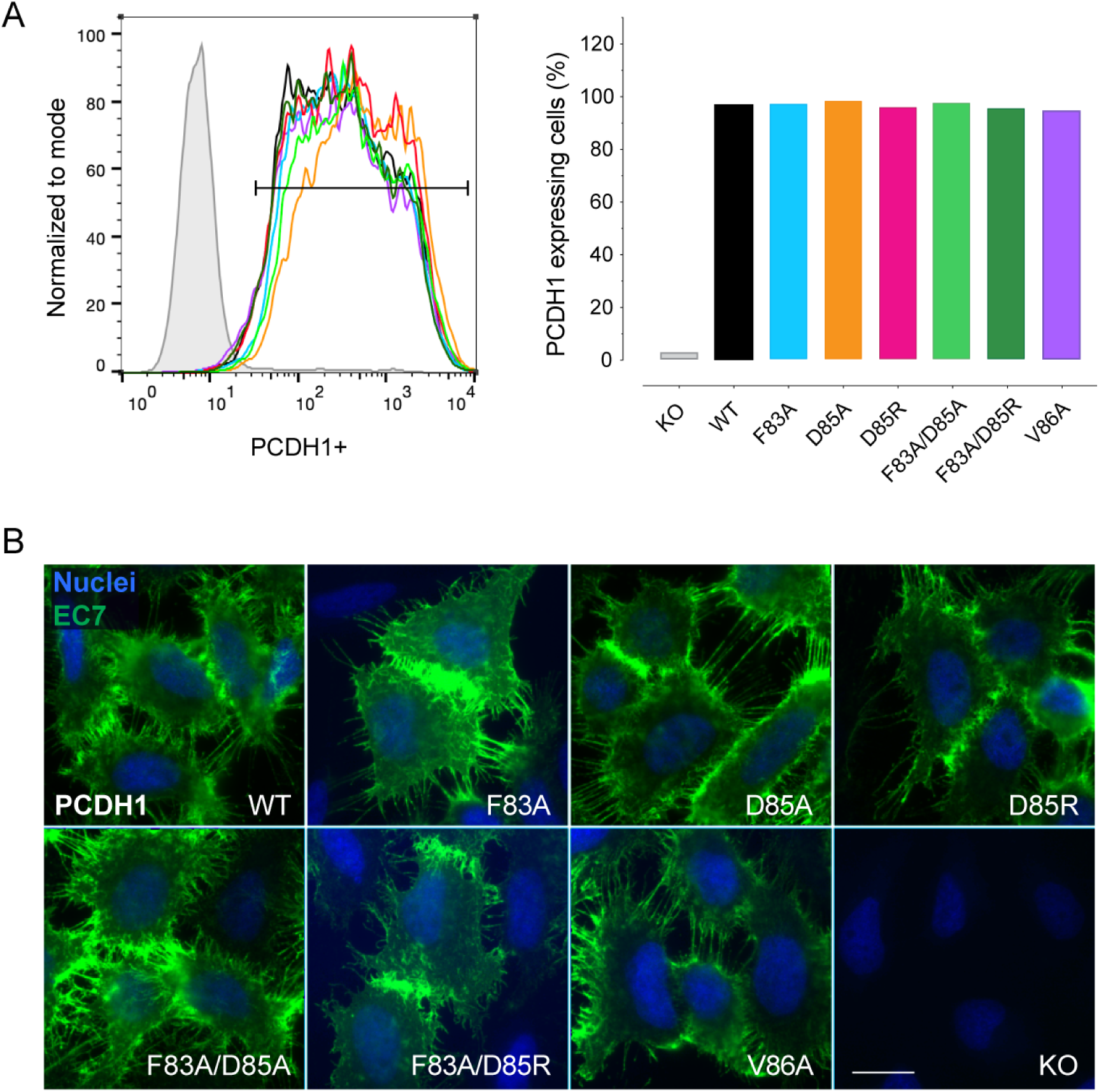
Overexpression of mutant PCDH1 exhibit similar expression levels to WT. (A) Flow cytometry histogram of PCDH1 expression and percent PCDH1^+^ cells (U2OS *PCDH1-*KO cells complemented with WT or mutant PCDH1). The cell surface-expressed PCDH1 protein was detected by an EC7-specific 3677-mAb and analyzed using flow cytometry. Data from one representative experiment is shown. (B) Surface expression of PCDH1 on cell lines from (A) visualized using an EC7-specific 3677-mAb. Scale bar, 20 µm. Representative images of two independent experiments are shown.

**Figure S6.**
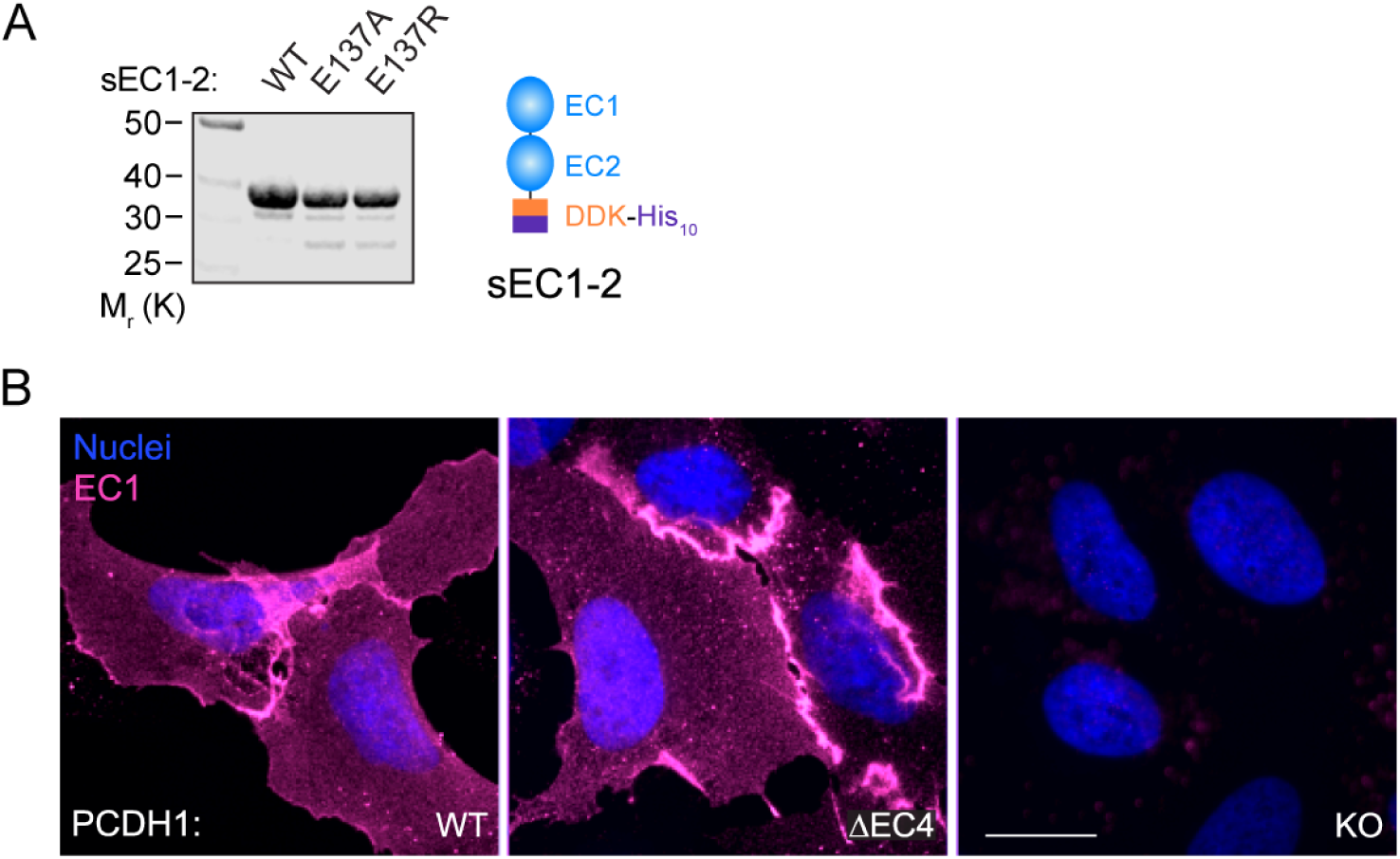
Mutations disrupting the EC1:EC4 interface in soluble PCDH1 and monomeric PCDH1 cell lines exhibit similar expression levels to WT. (A) Purified WT and mutant sEC1-2 were separated on an SDS-polyacrylamide gel and visualized by Coomassie Brilliant Blue staining. M_r_, relative molecular weight (K denotes x 1,000). Schematic representation of sEC1-2 (right). (B) Surface expression of PCDH1 on U2OS *PCDH1*-KO cells complemented with WT or mutant-without the EC4 domain (ΔEC4) PCDH1 proteins. The cell-surface, expressed PCDH1 was immunostained using a PCDH1 EC1-specific 3305-mAb. Scale bar, 20 µm. Representative images illustrating two independent experiments are shown.

**Figure S7.**
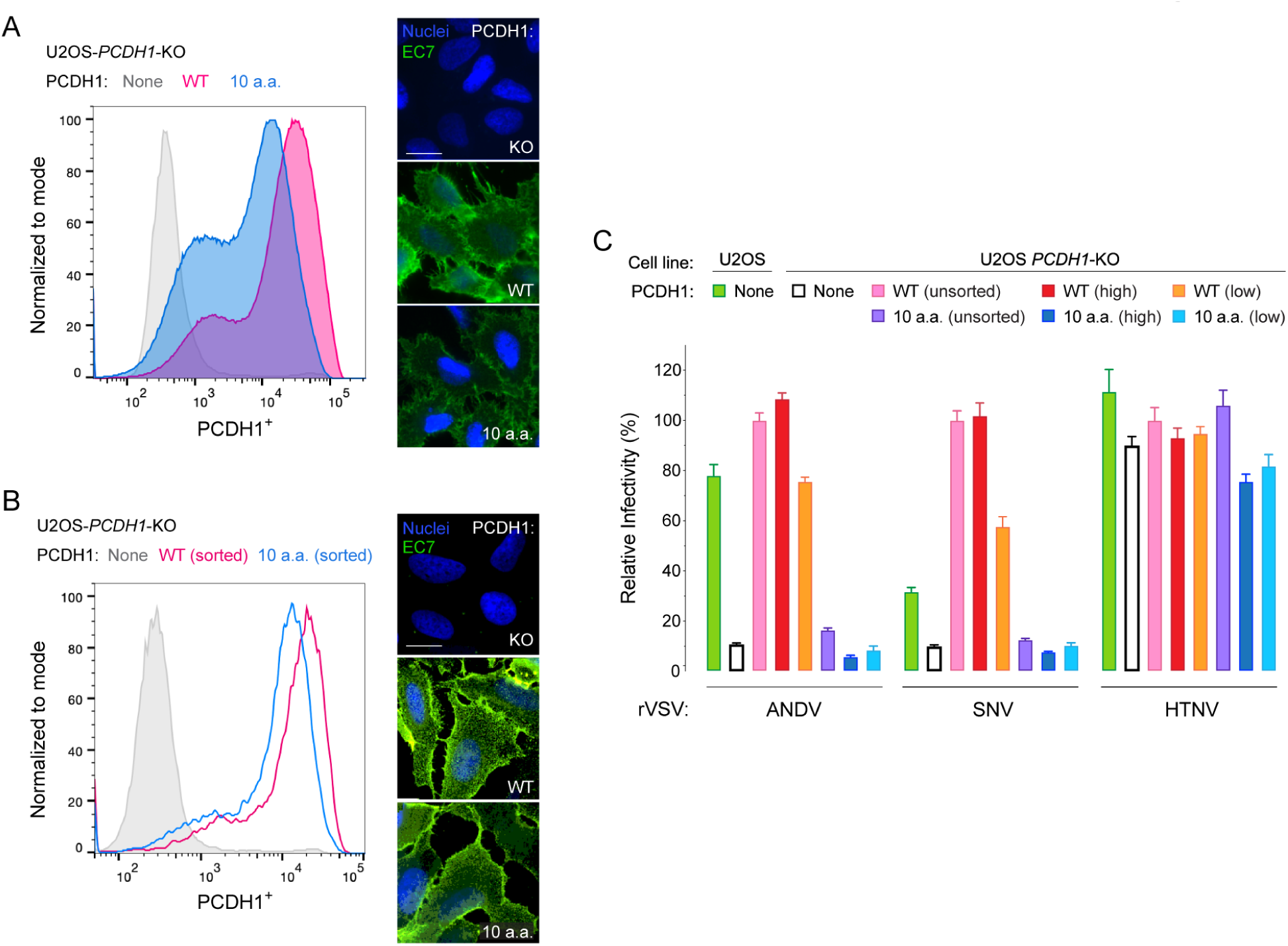
Cells expressing high levels of PCDH1(10a.a.) mutant do not support ANDV Gn/Gc-mediated infection. (A) Flow cytometry histogram and immunofluorescence of cell surface PCDH1 expression on U2OS *PCDH1*-KO cells complemented with either WT or mutant PCDH1 proteins. The surface of the cells was immunostained using a PCDH1 EC7-specific 3677-mAb followed by analysis using flow cytometry or imaged via an inverted fluorescence microscope. Representative flow cytometry histogram and representative fluorescent images, from two independent experiments, are shown. Scale bar, 20 µm. (B) Flow cytometry histogram and immunofluorescence images of sorted stably expressing cells, stained as described in (A). Cells described in (A) were sorted into high or low PCDH1 expression. Cells expressing high levels of PCDH1 are represented in the histogram and fluorescent images. Flow cytometry histogram from one experiment is represented and representative fluorescent images from two independent experiments are shown. Scale bars, 20 µm. (C) Relative infectivity of rVSVs bearing ANDV-, SNV-, or HTNV-Gn/Gc on U2OS cells, unsorted and sorted U2OS *PCDH1-*KO cells complemented with either WT or mutant PCDH1 described in (A) and (B). Averages ± SD: two to three experiments, *n* = 6-9.

**Figure S8.**
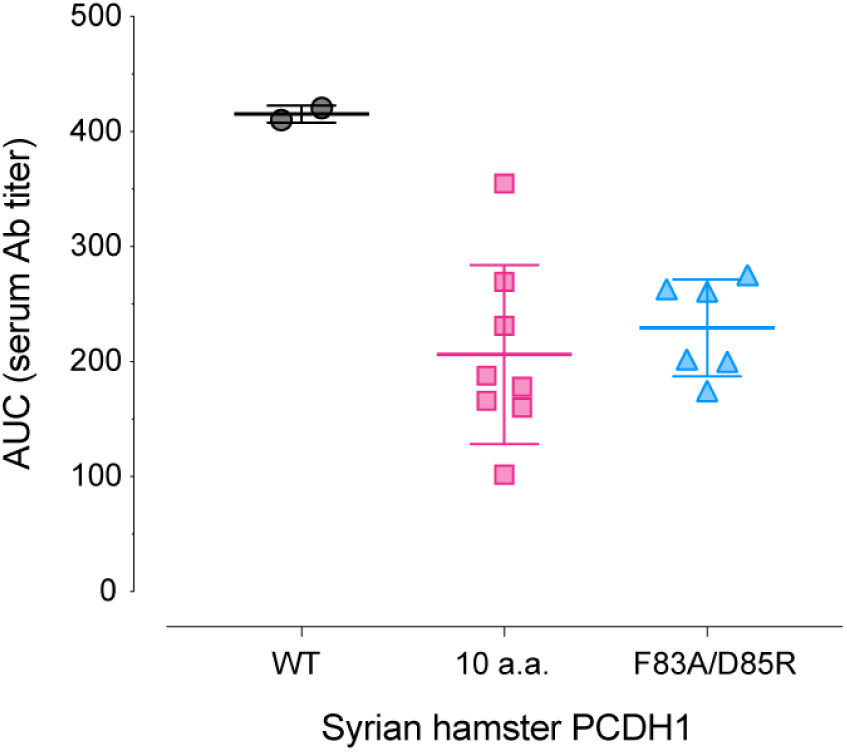
Seroconversion of Syrian hamsters challenged with ANDV. Area under the curve (AUC) for ANDV Gn/Gc-specific antibodies from WT, PCDH1(10a.a.), and PCDH1(F83A/D85R) CRISPR knock-in mutant Syrian hamsters’ sera. Syrian hamsters were challenged with 2,000 PFU of ANDV and serum titers were evaluated for all hamsters who survived until day 35. ANDV Gn/Gc-specific IgGs were detected using rVSV-ANDV-Gn/Gc coated ELISA plates. Averages ± SD; WT: *n* = 2, PCDH1(10 a.a.): *n* = 8, and PCDH1(F83A/D85R): *n* = 6.

